# Impairment of the cellulose degradation machinery enhances fungal virulence but limits reproductive fitness

**DOI:** 10.1101/2021.10.08.463612

**Authors:** Francisco M. Gámez-Arjona, Stefania Vitale, Aline Voxeur, Susanne Dora, Sascha Müller, Gloria Sancho-Andrés, Juan Carlos Montesinos, Antonio Di Pietro, Clara Sánchez-Rodríguez

**Author notes:** IPSP-Instituto per la Protezione Sostenibile delle Piante, CNR, 80055 Portici, Italy.

## Abstract

Fungal endophytes grow in the apoplastic space, in constant contact with the plant cell wall (CW) that hinders microbe progression, while representing a source of nutrients. Although numerous fungal CW modifying proteins have been identified, their role during host colonization remains underexplored. Here we show that the root-infecting plant pathogen *Fusarium oxysporum* (Fo) does not require its complete arsenal of cellulases to infect the host plant. Quite the opposite, Fo mutants impaired in cellulose degradation become hypervirulent by enhancing the secretion of virulence factors. On the other hand, the reduction on cellulase activity had a severe negative effect on saprophytic growth and microconidia production during the final stages of the Fo infection cycle. These findings enhance our understanding on the function of plant CW degradation on the outcome of host-microbe interactions and reveal an unexpected role of cellulose degradation in a pathogen’s evolutionary success.

**Teaser:** Unexpectedly, fungi compromised in their capacity to degrade plant cellulose are hypervirulent but impaired in sporulation.

## Introduction

Most plant-microbe interactions are initially established at the apoplast. Here, the plant CW constitutes a source of nutrients and a physical barrier for the intruder, making its modification an important aspect of microbe colonization of the host. The main components of plant CWs are the para-crystalline fibres of cellulose embedded in a matrix of hemicellulose, pectin, lignin and suberin (*1*–*3*). Ions and wall remodeling proteins modify the interactions between CW polysaccharides *in muro* changing the physicochemical properties and, consequently, the structure and function of the CW (*4, 5*). The CW composition is not homogenous among different plant species, tissues or even within a single cell (*6*). This molecular complexity forces plant-colonizing microbes to metabolically regulate the secretion of a broad set of CW modifying proteins (CWMP) required to loosen and digest the plant CWs they encounter (*7, 8*). The majority of these CWMPs are CW degrading enzymes (CWDEs), including glycosyl-hydrolases, lytic polysaccharide monooxygenases, pectate and pectin lyases, and esterases, which are further divided into different subgroups based on sequence similarity in the catalytic domain (*9*–*13*). Although all plant-colonizing microbes encode for CWMPs, it has been suggested that the role of these enzymes is more prominent in pathogens, which rely on their activity during host infection (*14*–*18*). However, so far the large functional redundancy of CWDEs has largely prevented a complete assessment of their role (*19*), since deletion of individual CWMEs genes do not generally affect the virulence of the pathogen (*20*). Fungi have evolved transcription factors that co-regulate the expression of CWMP groups based on the composition of their environment. Therefore, it has been suggested that targeting such master transcriptional regulators should allow the assessment of the role of a given group of CWMPs (*21*).

In their co-evolution journey, while microbes adapted to loosen and break the host CWs, plants evolved to perceive these degradation products as damage-associated molecular patterns (DAMPs). These DAMPs, together with microbe-associated molecular patterns (MAMPs) such as fungal chitin (*22*), activate the plant pattern-triggered immunity against the intruder (*23, 24*). The best characterized CW-derived DAMPs are pectin-derived oligogalacturonides (OGs), resulting from pectinase activity (*25*–*28*). Cellulose degradation products, i.e., like cellobiose, have also been shown to induce plant defense and are, therefore, considered potential DAMPs, although their release upon microbe colonization has not yet been proven (*25*–*27*).

The plant CW characteristics directly influence the communication between the two organisms. This is particularly relevant for plant response to microbes that mainly live in the apoplast, such as vascular fungi belonging to the *Fusarium oxysporum* (Fo) species complex, which are further classified into different formae speciales (ff.spp.) based on their host preference (*29*). Fo_s_ are soil-borne microbes that colonize the roots of many plant species, being responsible for the devastation of many economically important crops throughout the world (*29*). Fo_s_ are considered pathogenic when they cause plants to wilt and eventually die, which occurs because water flow and nutrient uptake are blocked by fungal proliferation inside the xylem. To reach the vascular system, Fo grows predominantly in the root apoplast (*30, 31*), making this fungus an ideal model to study CW-microbe interaction.

Cellulose is the most abundant and most recalcitrant component of the CW that provides mechanical strength to the plant cells thanks to its paracrystalline structure (*7*). Consequently, its synthesis is tightly regulated in response to intruders, including Fo (*32*–*34*) cellulose is one of the main targets of microbial CWMPs. Pathogens secrete a broad range of cellulolytic enzymes (glycosyl-hydrolases (GHs) and lytic polysaccharide monooxygenases (AAs)) (*16*), suggesting that this family of enzymes should be essential for infection. An optimal degradation of complex carbohydrates requires a hierarchical metabolic response. In the case of crystalline cellulose, AAs disrupt the outermost crystalline part of the cellulose fiber, thereby allowing the remaining cellulases to participate in the degradation process (*9, 10, 35*). The transcriptional regulation of fungal cellulases has been reported to be dependent on a conserved zinc binuclear cluster transcription factor named CLR1. CLR1 regulates the expression of cellulolytic enzymes in *Neurospora crassa* and other ascomycetes and is, therefore, required for growth on cellulose as the sole carbon source (*7, 36, 37*).

Here we investigated the impact of cellulose degradation during microbe infection by identifying and targeting the orthologues of *N. crassa CLR1* in Fo f.sp. *conglutinans* (Fo5176) and *lycopersici* (Fol4287), pathogens of *Arabidopsis thaliana* and *Solanum lycopersicum*, respectively (*38, 39*). Unexpectedly, our data show that crystalline cellulose degradation is not only dispensable, but disadvantageous, for Fo infection, as shown by the increased virulence displayed by *clr1* mutants. Using the model pathosystem *Arabidopsis thaliana*-Fo5176 ((*33, 34, 40, 41*), we show that the *clr1* mutant compensates for cellulose degradation deficiency by increasing the secretion of various virulence factors to compromise plant immune responses. On the other hand, the reduction of cellulose degradation capacity severely compromised fungal metabolism during its saprophytic growth, with dramatic consequences for its microconidia production. Taken together, our findings expand the current understanding of plant-microbe interactions, showing that the cellulose is not the assumed central physical barrier against fungi, but a primary source of carbon to complete the pathogen’s life cycle in the plant.

## Results

### The transcriptional regulator CLR1 is required for efficient cellulose degradation by Fo5176

To understand the role of Fo cellulases in root infection, we aimed to obtain a Fo5176 mutant impaired in cellulose degradation. Considering the large number of putative cellulases expressed by Fo5176 during root colonization (*34*) and their high functional redundancy, we chose to target the transcription factor CLR1, a master regulator of fungal cellulase expression previously identified in *N. crassa* (*36*). Using an *in silico* analysis, we found CLR1 orthologues in various Fusarium genomes, sharing a protein sequence identity above 70% and having the same DNA-binding domain sequence (Fig. 1A and 1B). Therefore, we generated a CLR1 null mutant in the previously described Fo5176 *pSIX1::GFP* background, which allows for monitoring Fo5176 colonization of the root vasculature (*33, 41*). We obtained deletion mutants lacking the entire *CLR1* coding region (*clr1-1* and *clr1-2)* and a complemented strain (*clr1C)* generated by reintroducing the wild-type *pCLR1::CLR1* into the *clr1-1* mutant background (Fig. S1). We first determined whether *clr1* mutants were affected in stress response by growing them in control conditions (YPD), salt stress (NaCl), osmotic stress (Sorbitol), CW stress (Congo Red and Calcofluor) (*42*) as well as on minimal medium. There were no detectable differences in the colony phenotype of *clr1* mutants compared to the wild type (WT) in any of these conditions (Fig. S2). Next, we evaluated the role of CLR1 in cellulose degradation by Fo by testing the ability of *clr1-1, clr1C* and WT to utilize different carbon sources such as sucrose, cellobiose and cellulose. Fungal growth on sucrose was similar across all genotypes (Fig. 1C and S3). The *clr1-1* mutant exhibited a significant growth reduction on cellobiose and only showed residual growth on cellulose compared to WT or *clr1C* (Fig. 1C, 1D and S3). A similar phenotype was observed for *clr1-2* (Fig. S4A). These data are similar to those described in *N. crassa* (*36*), indicating that the essential function of CLR1 in cellulose catabolism is conserved across different ascomycetes.

**Fig. 1.**
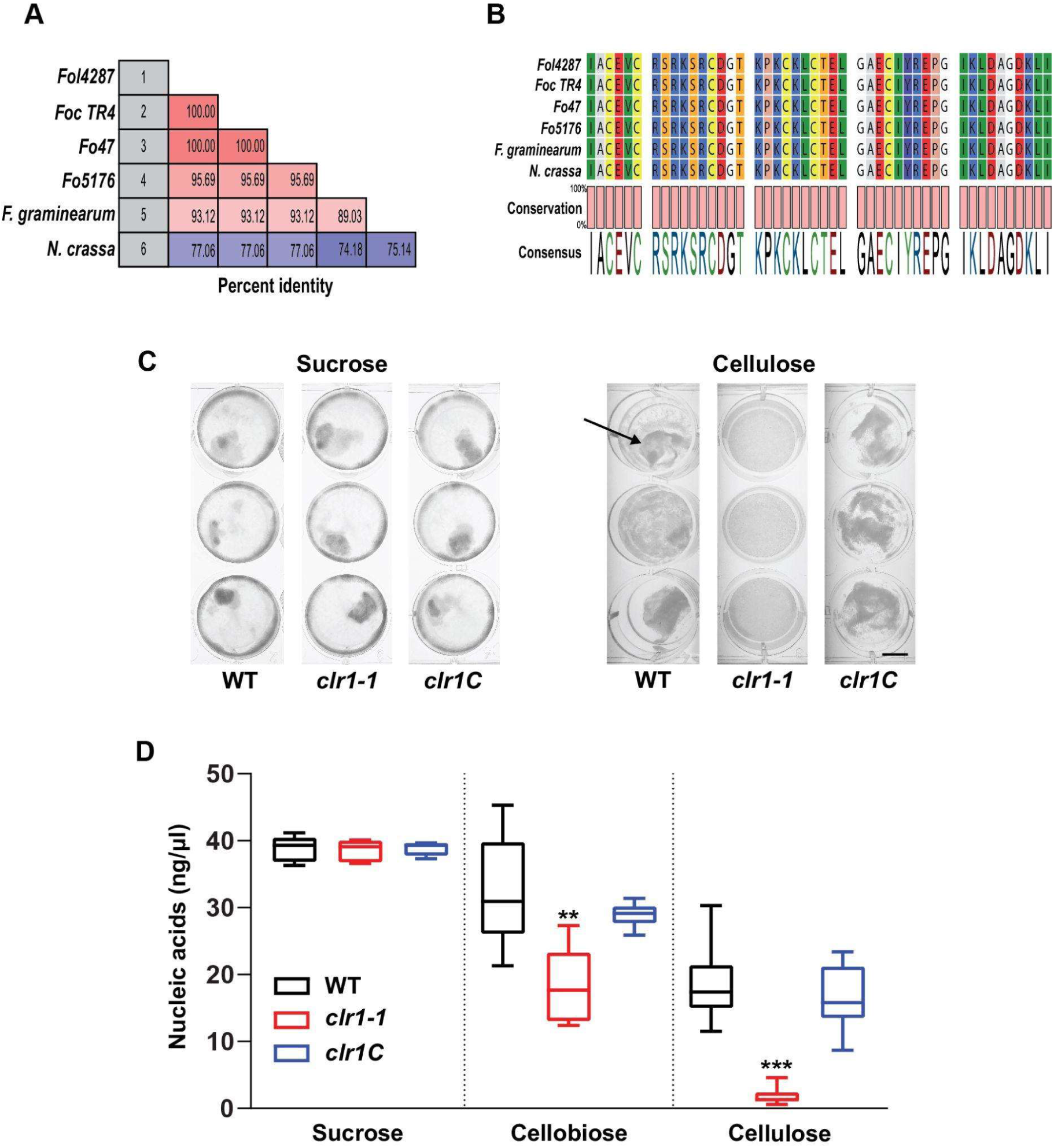
Cellulose degradation is regulated by CLR1 in *F. oxysporum* (Fo5176). **(A)** Pairwise comparison of 5 Fusarium species or Fo species complexes to identify potential orthologous to the *Neurospora crassa* transcription factor. The percent of protein identity is indicated. **(B)** Sequence alignment of the DNA-binding domains of the CLR1 proteins identified in (A) The conservation grade is shown. Note the high conservation of the Zn(2)-C6 fungal-type domain, ACEVCRSRKSRCDGTKPKCKLCTELGAECIY, related to the DNA-binding domain. **(C)** Representative picture of WT, *clr1-1* and *clr1C* growth on sucrose (left panel) or cellulose (right panel). The arrow indicates the presence of mycelia. Scale bar, 1cm. **(D)** Growth of WT, *clr1-1* and *clr1C* on different carbon sources measured as nucleic acid concentration (ng/μl). The strains were growing on sucrose 0.5% or cellobiose 0.5% for 3 days, or on cellulose 0.5% for 7 days. Shown are the box plots: centerlines show the medians; box limits indicate the 25^th^ and 75^th^ percentiles; whiskers extend to the minimum and maximum, n≥5. Asterisk indicated differences relative to WT. Welch’s unpaired t-test; *** p-value<0.001. ** p-value ≤ 0.01.

### Loss of CLR1 leads to faster colonization of the root cortex and xylem

To determine whether Fo requires cellulose degradation to infect its plant hosts, we performed plate infection assays as described before (*41*). We monitored root vascular colonization by following GFP signal presence in the root vasculature at different days post-transfer (dpt) to spore-containing plates. Unexpectedly, we counted a significantly higher number of vascular penetrations of *clr1-1* than WT at all time points, correlated with a higher expression of *FoSIX1* and an increased *in planta* fungal biomass at 7dpt measured by qRT-PCR, which was restored to WT levels in the *clr1C* straine (Fig. 2A-C). A second isogenic mutant, *clr1-2*, also showed a higher capacity than WT to reach the host xylem (Fig. S4B). We therefore focused on the *clr1-1* mutant for further characterization. Using confocal microscopy, we observed that *clr1-1* advanced faster than WT through the epidermal apoplast as could be seen in the epidermis-cortex interface at 3dpt with a significant higher frequency than the WT fungus (Fig. 2D and 2F). These results indicate that *clr1-1* is more efficient than WT in growing through the host CWs and reaching the xylem and might explain the higher levels of vascular invasion observed at later stages of the infection (Fig. 2A-C).

**Fig. 2.**
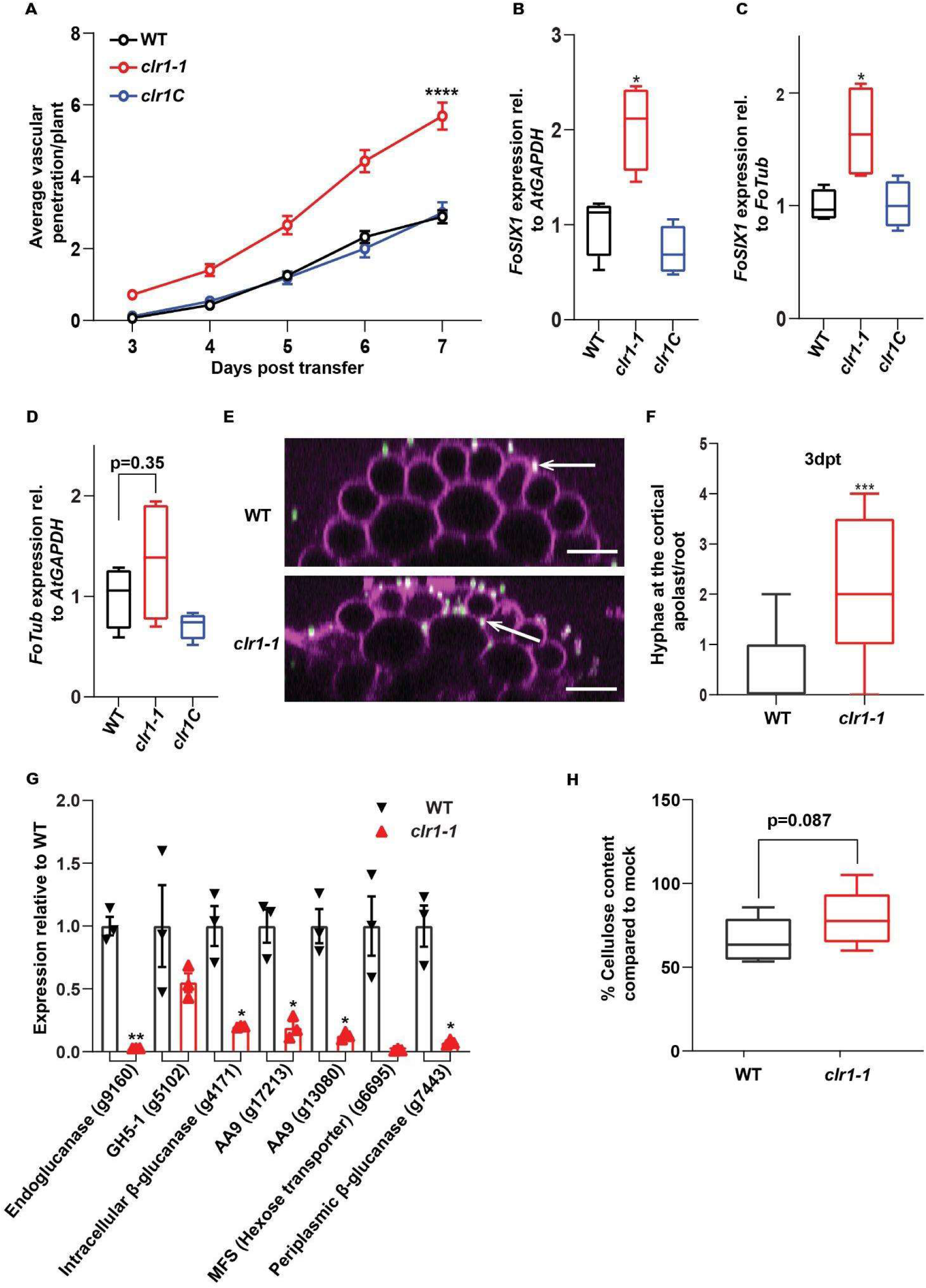
The lack of CLR1 increases Fo pathogenicity. **(A)** Cumulative Arabidopsis root vascular penetration by Fo at different days post-transfer (dpt) to WT, *clr1-1* or *clr1C* microconidia-containing plates. Values are mean +/- SEM, N≥ 28 plants from one representative experiment. The experiment was performed 3 times with similar results. RM two-way ANOVA on vascular penetration rate: p<0.0001 (fungal genotype), p<0.0001 (time), p≤0.0001 (fungal genotype x time). Asterisk indicated a statistical difference with respect to WT at 7 dpt, Tukey’s multiple comparison test, **** p<0.0001. **(B)** and **(C)** *FoSIX1* expression relative to *AtGAPDH* or *FoTub*, respectively, in infected Arabidopsis roots at 7 dpt as in (A).. Shown are the box plots: centerlines show the medians; box limits indicate the 25^th^ and 75^th^ percentiles; whiskers extend to the minimum and maximum. N=4 biological replicates; Welch’s unpaired t-test; * p-value ≤0.05. **(D)** Fungal biomass calculated as *FoTub* expression relative to *AtGAPDH*. Box plots shown as above. N=4 biological replicates. **(E)** Representative confocal image of WT or clr1-1 hyphae (green) colonization of Arabidopsis roots stained with PI (magenta). Arrows point to hyphae. Scale bars: 20μm. **(F)** Number of WT or *clr1-1* hyphae able to reach the cortical apoplast as shown in (E). Data are represented on box plots as described above. N≥12, Welch’s unpaired t-test; *** p-value<0.001. **(G)** WT and *clr1-1* gene expression in 7 dpt infected roots normalized to the fungal *FoTub* gene. Bars represent means +/- SEM, N=3 biological replicates (arrowheads). For each gene, the data were normalized to WT; Welch’s unpaired t-test; * p-value<0.05. ** p-value<0.01. **(H)** Cellulose content in WT and *clr1-1* infected roots at 7dpt represented as % of cellulose measured in 7 dpt mock roots. Shown are the box plots as described above. N=8 biological replicates. Welch’s unpaired t-test; p-value indicated.

At 7dpt, when the fungus is colonizing the vasculature in all roots, we observed a reduced expression of genes required for cellulose degradation in *clr1-1* compared to WT including endoglucanase, glycosyl hydrolase family 5 member, two AA9 enzymes, and a periplasmic □-glucanase, as well as for sugar metabolism such as a plasma membrane hexose transporter and an intracellular □-glucanase (Fig 2G). These are orthologues of *N. crassa* genes previously reported to be regulated by CLR1 during growth on cellulose (*43*). Importantly, we detected a lower amount of residual cellulose in roots infected with WT compared to those infected with *clr1* (Fig. 2H), in line with the results of growth on cellulose (Fig. 1C). Taken together, our data reveal that CLR1 is required by Fo for efficient degradation of cellulose during root colonization but dispensable for reaching the plant vasculature.

### Arabidopsis defense response is delayed upon *clr1-1* infection compared to WT

The reduction in cellulase activity observed in *clr1-1* might reduce the amount and/or nature of the CW-derived DAMPs generated by this mutant compared to the WTs. Therefore, the increased virulence of the cellulase-deficient *clr1* mutant might be a consequence of an impaire activation of plant immune responses. To address this hypothesis, we measured the expression of three genes previously reported to be activated in Arabidopsis roots infected by Fo5176; *At1g51890, WKRY45* and *WKRY53 (33, 44)*. At 2 and 3dpt, the expression of the three defense genes was lower in plants infected with *clr1* than WT (Fig. 3A). These differences progressively disappeared with the increased vascular colonization. At 7dpt, when the xylem was abundantly occupied by *clr1-1*, the plant defense gene expression was higher than in roots infected with WT (Fig. 3A). These findings suggested that the higher number of vascular penetrations observed upon *clr1-1* infection (Fig. 2A) might be a consequence of delayed activation of defense related genes during the early stages of the interaction.

**Fig. 3.**
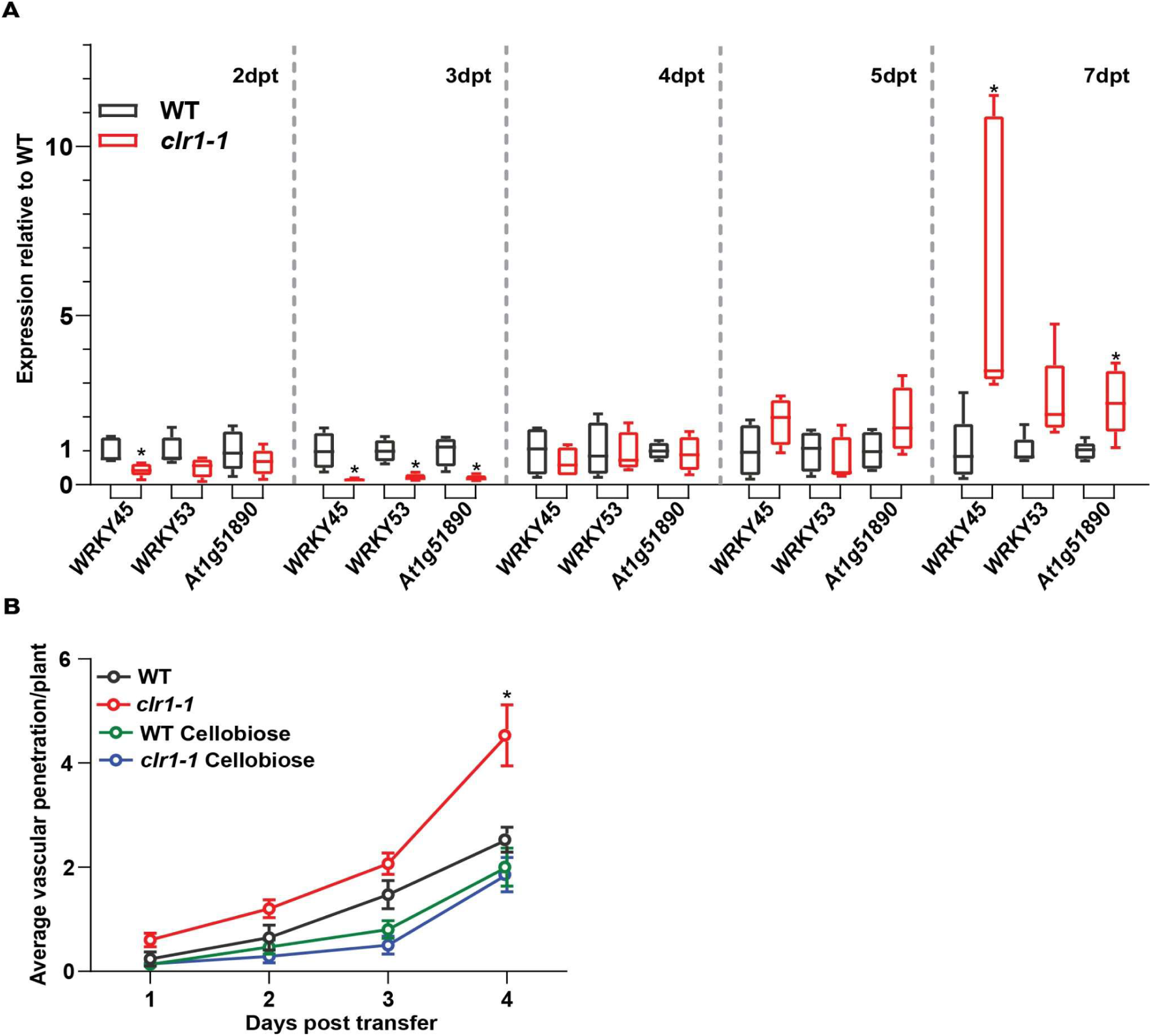
The plant perceives later *clr1-1* than WT. **(A)** Expression of *WRKY45, WRKY53*, and *At1g51890* relative to *AtGAPDH* in Arabidopsis roots at 2, 3, 4, 5 or 7dpt to plates with WT or *clr1-1* microconidia. Shown are box plots: centerlines show the medians; box limits indicate the 25^th^ and 75^th^ percentiles; whiskers extend to the minimum and maximum, n≥4 biological replicates. For each gene, the data were normalized to WT and the statistical differences are indicated by asterisks; Welch’s unpaired *t*-test; * p-value<0.05. **(B)** Cumulative Arabidopsis root vascular penetration by Fo at different dpt to WT, *clr1-1* or *clr1C* hyphae-containing plates. Half of the plants were pretreated with cellobiose for 25 minutes before being exposed to the fungus. Values are mean +/- SEM, N≥14 from one representative experiment. The experiment was repeated three times with similar results. RM two-way ANOVA on vascular penetration rate: p<0.0001 (fungal genotype and treatment), p<0.0001 (time), p≤0.0001 (fungal genotype x time). Asterisk indicates a statistical difference with respect to WT at 7 dpt. Tukey’s multiple comparisons test, * p<0.05.

We next tested whether external addition of CW-derived DAMPs could reducere the virulence of *clr1-1* to WT levels. We used cellobiose, a cellulose-degradation product confirmed to induce plant defense (*27*), to prime the root immune system. We pretreated plants for 25 minutes with this molecule before transferring them to plates containing 2-day-old WT or *clr1-1* mycelium. We confirmed that cellobiose-induced root-defense priming returned the plant susceptibility of *clr1* to WT levels (Fig. 3B).

### Different amounts of CW degradation products accumulate upon *clr1-1* infection

The reduced cellulose degradation and the concomitant absence or decrease of cellobiose in the apoplast during *clr1-1* infection might explain the delay in perception of this mutant by the root and its increased virulence t. To test this idea, we measured the oligosaccharides derived from plant CW degradation by Fo5176 using HP-SEC-MS/MS (*45*). By employing different commercial standards to identify the different hexoses and saturated OGs (Fig. 4A), we identified cellobiose in Fo5176-infected but not mock-treated roots confirming its predicted role as DAMP. Our results showed a shift in the di-hexose profiles from sucrose-to cellobiose-enriched at 2dpt and 3dpt, respectively (Fig. 4B). From 3 dpt on the amount of di-hexoses, mainly cellobiose, was higher during infection with *clr1-1* compared with then WT (Fig. 4B and S5). The HP-SEC-MS results indicate that the impaired capacity of the root to detect *clr1-1* can not be explained by a reduced release of cellobiose by the mutant. Therefore, we studied the OGs produced by Fo5176 and identified acetylated OGs as a main product of plant CW degradation by both *clr1* and WTi (Fig. 4B, 4D, 4E and S5). Moreover, less OGs were detected during *clr1* infection compared to WT-infected roots, especially at 3 and 4dpt (Fig 4C-E and S5). This difference was statistically significant for Gal_4_Ac and Gal_2_ox (Fig. 4E and 4D). The decreased accumulation of OGs during the early stages of plant infection is in line with the compromised perception of *clr1-1* by the root. We detected significant differences in the amount of plant CW-degradation products in WT- and *clr1-1* infected roots, that might act as DAMPs. Therefore, we next tested whether WT-generated DAMPs can restore plant susceptibility to *clr1-1* by performingplate infection assays using the same number of microconidia concentration of the *clr1-1*, WT or both (50% WT+50% *clr1-1*). We expected to see a decrease in *clr1-1* virulence proportional to the concentration of WT spores in the mixed treatment if DAMP accumulation is the main reason for the differential plant susceptibility to *clr1-1* and WT6. By contrast, we found that the 50%-mixed infection displayed the same pathogenicity as *clr1-1* alone (Fig. S6A), observing a decrease in xylem penetration only when 90% of the spores were WT (Fig. S6B). Our data indicate that although *clr1-1* infection generates different amounts of CW-degradation products than WT, they alone do not explain *clr1-1* higher pathogenicity.

**Fig. 4.**
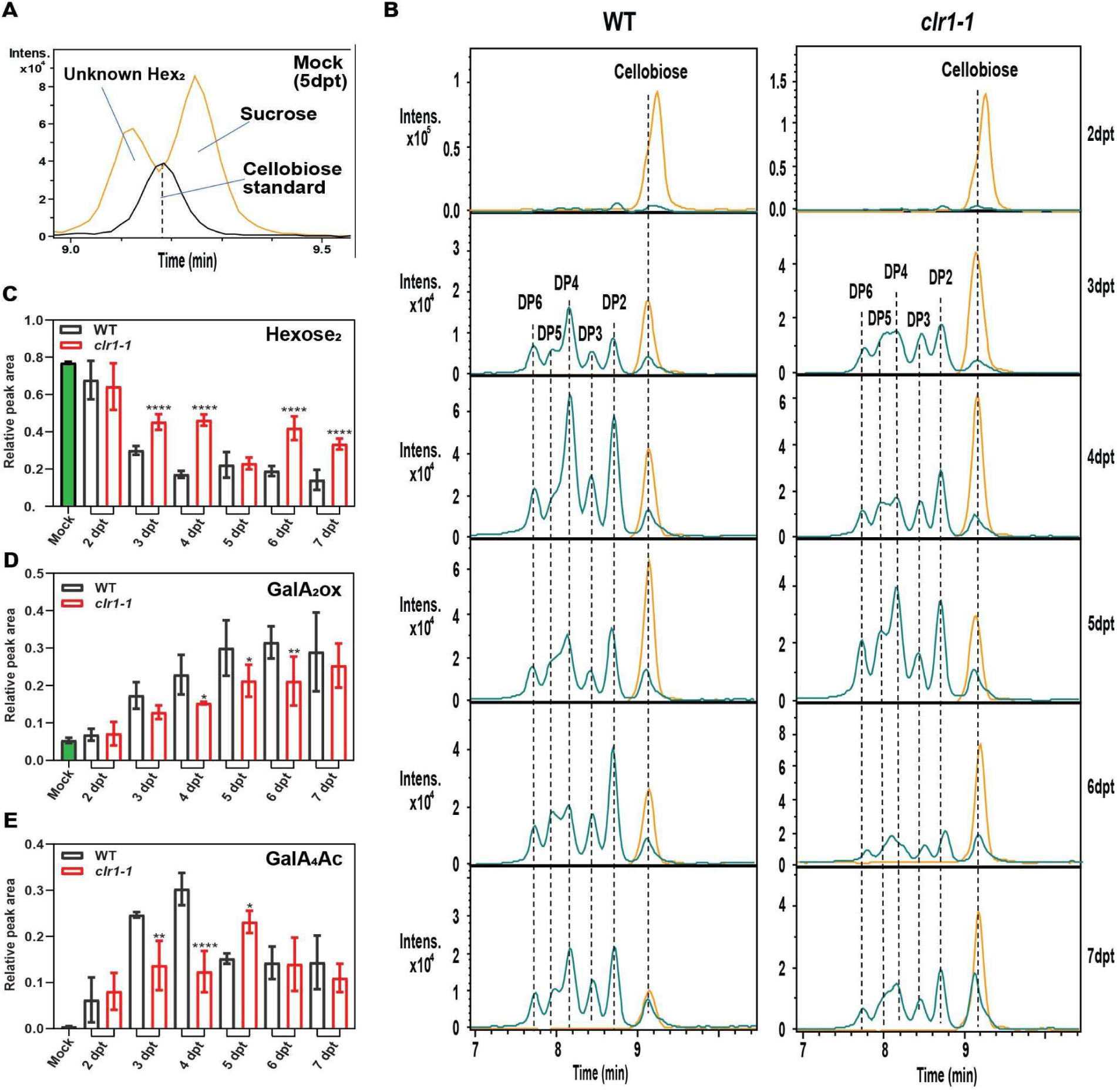
*clr1-1* infection generates different amounts of plant cell wall-derived molecules than WT. **(A)** Extracted ion chromatograms for m/z 341 of mock root cell wall (yellow) water extract (5 dpt) and cellobiose comercial standard (black) analysed by HP-SEC-MS. Saccharose and another di-hexose but no cellobiose are detected in cell wall water extract. **(B)** Representative extracted ion chromatograms of oligogalacturonides (OGs; blue) and di-hexoses (yellow) detected along 5 dpt to plates with WT or *clr1-1* microconidia. DP indicates the degree of polymerization of OGs. The retention time of different standards of DP and cellobiose is indicated with a vertical dotted line. **(C-E)** Kinetics of Hexose_2_ (C), GalA_2_ox (D), and GalA_4_Ac (E) produced in Arabidopsis roots by WT or *clr1-1* infection. Mock indicates the levels of polysaccharides at 5 dpt. Bars are means +/- SEM, N=3 biological replicates as described in (B) and 2 replicates for mock samples. Asterisk indicates statistical significance compared to the WT on each dpt, two-way ANOVA with LSD Fisher test post hoc comparison; **** p<0.0001** p<0.01, * p<0.05. OGs are named GalA_x_,GalA_x_Ac_y_ or GalAox with the subscript numbers indicating the degree of polymerization (x) and the number of acetylated groups (y); “ox” indicate the presence of oxidized groups.

### *clr1-1* secretes more virulence factors than WT during root colonization

Pathogenicity relies on the ability of microorganisms to evade the MAMP- and DAMP-induced host defenses using secreted and cell-surface localised virulence factors (*46*). We performed proteomic analysis of the secretomes of WT and *clr1-1* during plant infection to determine whether differences in their profile of secreted virulence factors could explain the higher virulencey of *clr1-1*. Seedlings growing in hydroponics were inoculated with either WT or *clr1-1* microconidia. 362 fungal proteins were identified, 75% with predicted secretory signal peptides. Of these, more than 50% belong to groups considered as virulence factors in other plant pathogens such as *F. graminearum* (*46*), including glycosyl hydrolases (26%), peptidases (13%), redox-related proteins (10%) and pectin and pectate lyases (4%) (Fig. 5A).

**Fig. 5.**
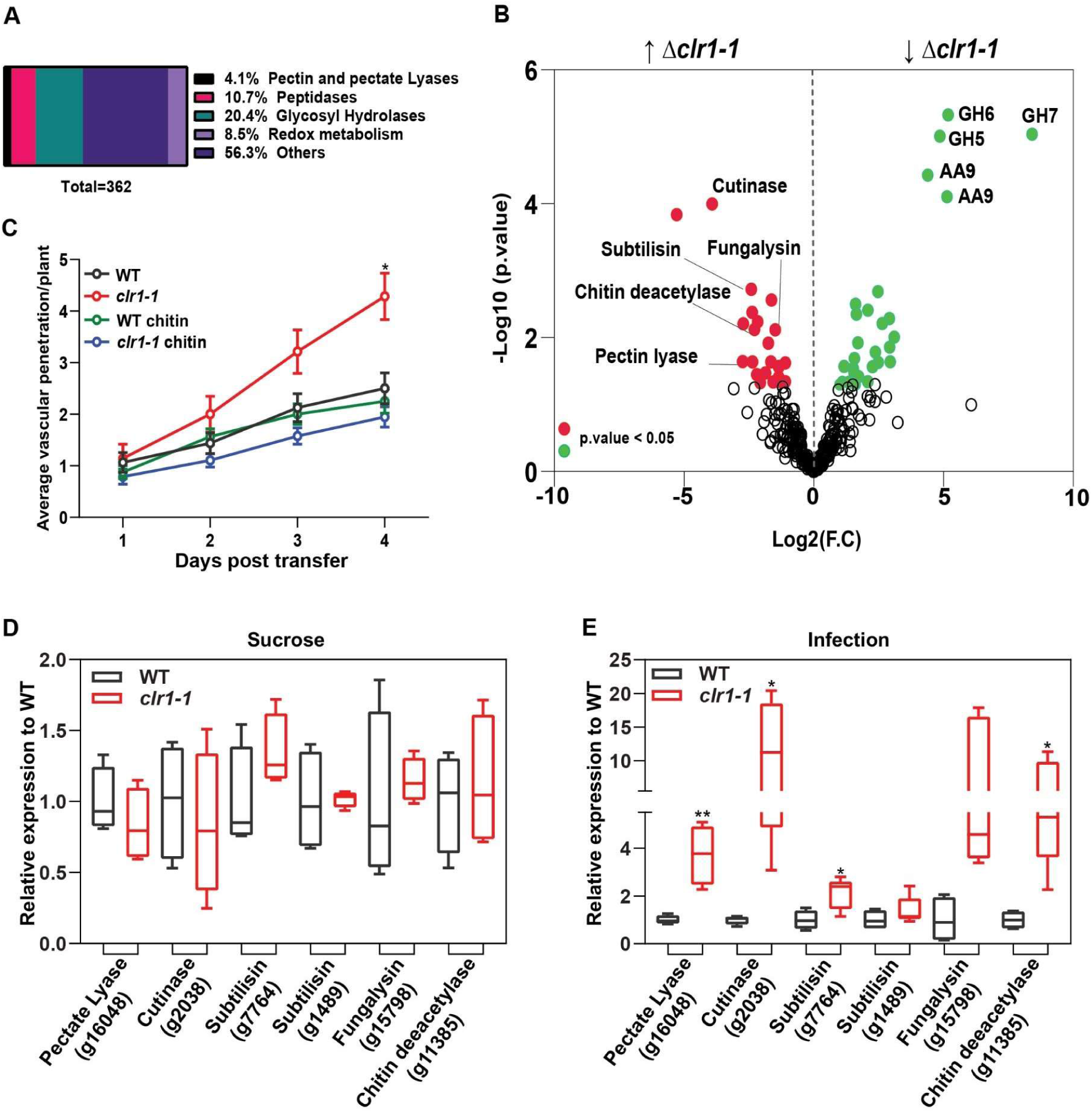
*clr1-1* secretes more virulence factors than WT during Arabidopsis root infection. **(A)** Proteins identified in the secretome of WT Fo at 3 dpt of Arabidopsis seedlings in hydroponic conditions. A total of 362 fungal proteins were identified in 4 independent replicates and the percentages of the most representative family proteins found are presented. **(B)** Volcano plot of differences in the abundance of proteins identified in the secretome of *clr1-1* relative to the WT. Proteins significantly (*moderated T-test, p*-values < 0.05) less or more present in the *clr1-1* secreteome are shown in green or red, respectively. The name or family of the most relevant proteins is indicated. **(C)** Cumulative Arabidopsis root vascular penetration by Fo at different dpt to WT, *clr1-1* or *clr1C* hyphae-containing plates. Half of the plants were pretreated with chitin for 25 minutes. Values are mean +/- SEM, N≥14 from one representative experiment. The experiment was repeated three times with similar results. RM two-way ANOVA on vascular penetration rate: p<0.001 (fungal genotype and treatment), p<0.0001 (time), p ≤ 0.0001 (fungal genotype x time). Asterisk indicates a statistical difference with respect to WT at 7 dpt. Tukey’s multiple comparisons test, * p<0.05. **(D)** and **(E)** Gene expression of 6 virulence factors relative to *FoTub* in WT or *clr1-1* grown 3 days in sucrose (D) or at 3dpt in Arabidopsis roots (E). The genes were selected among the ones enriched in the *clr1-1* secretome as shown in (B: : a pectate lyase (g16048), a cutinase (g2038), two subtilisins (g7764 and g1489), a fungalysin (g15798) and chitin deacetylase (g11385) Shown are box plots: center lines show the medians; box limits indicate the 25th and 75th percentiles; whiskers extend to the minimum and maximum, n ≥ 4. For each gene, the data were normalized to WT and the statistical differences are indicated by asterisks. Welch’s unpaired *t*-test; * p-value<0.05. ** p-value<0.01.

As expected, we found a significant reduction in the abundance of proteins required for cellulose degradation in the *clr1-1* secretome (Fig. 5B and Table 1), including GH5 (g17181), GH6 (g15944), and GH7 (g873) which are predicted to exhibit cellulolytic activity (*47*), as well as AA9s that make crystalline cellulose accessible to cellulases (*10*). These data confirm the downregulation of cellulase-related genes observed in *clr1-1* and explain its reduced capacity to degrade cellulose (Fig. 1C, 2D, 2F, 2G, and S3).

**Table 1.**
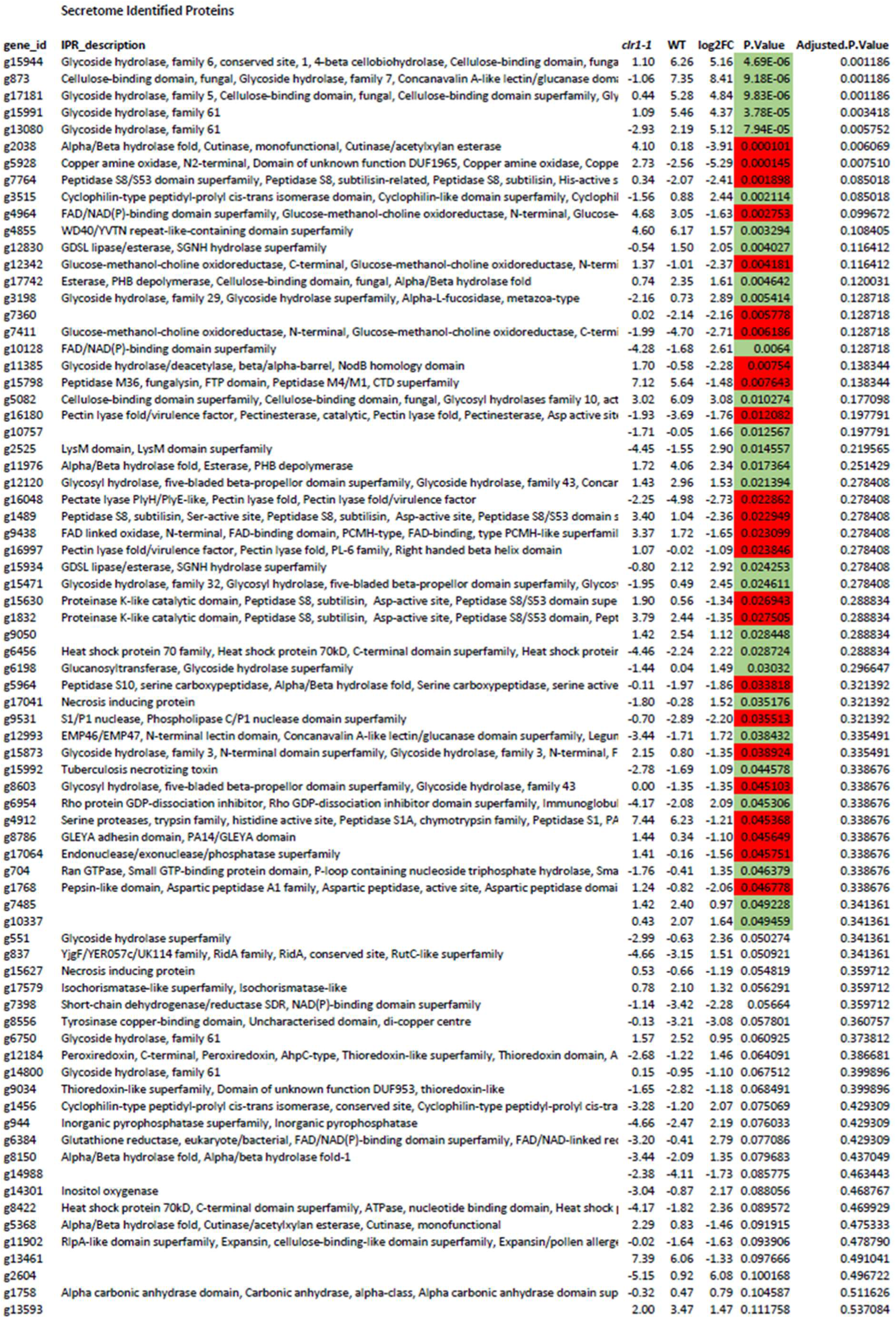
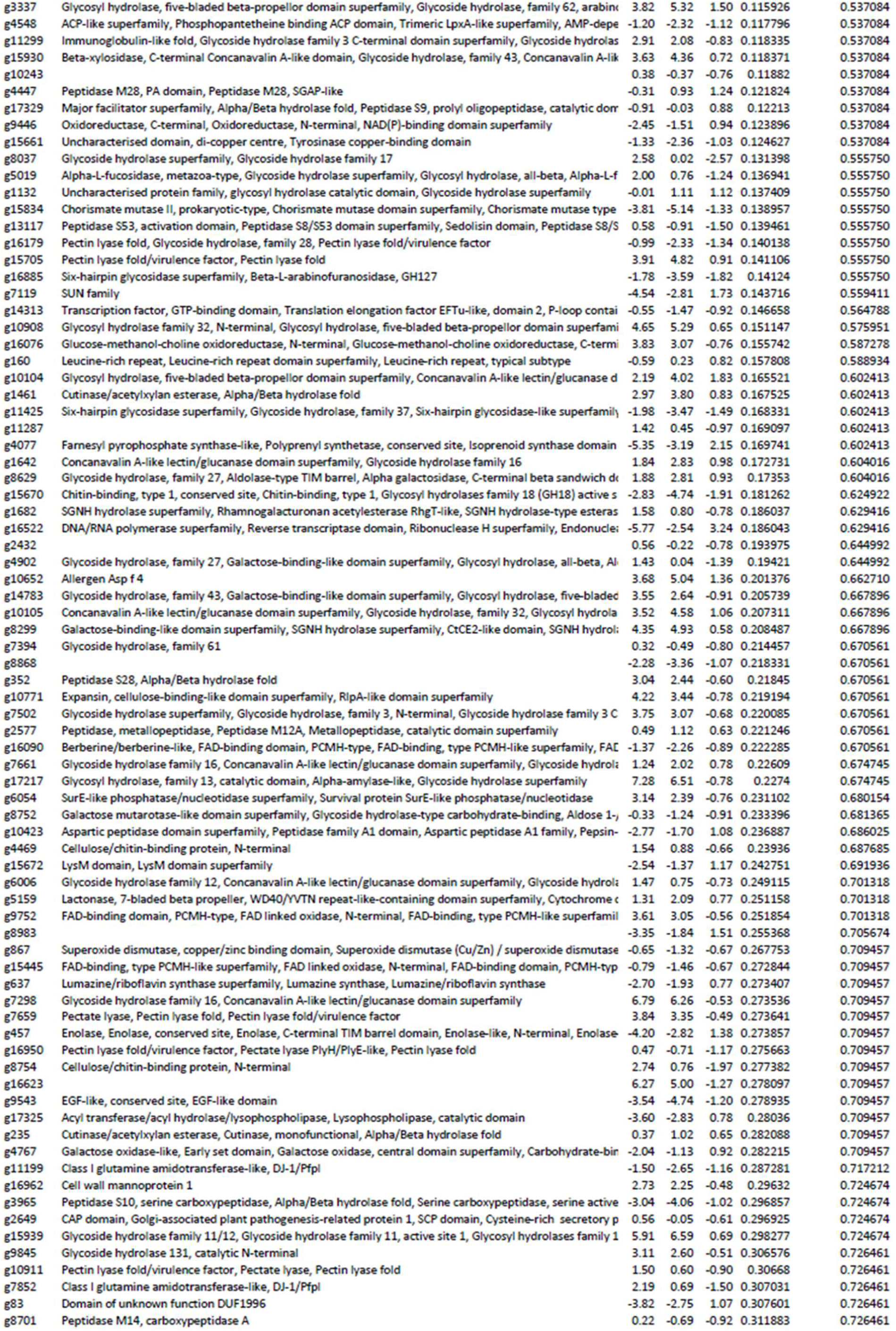
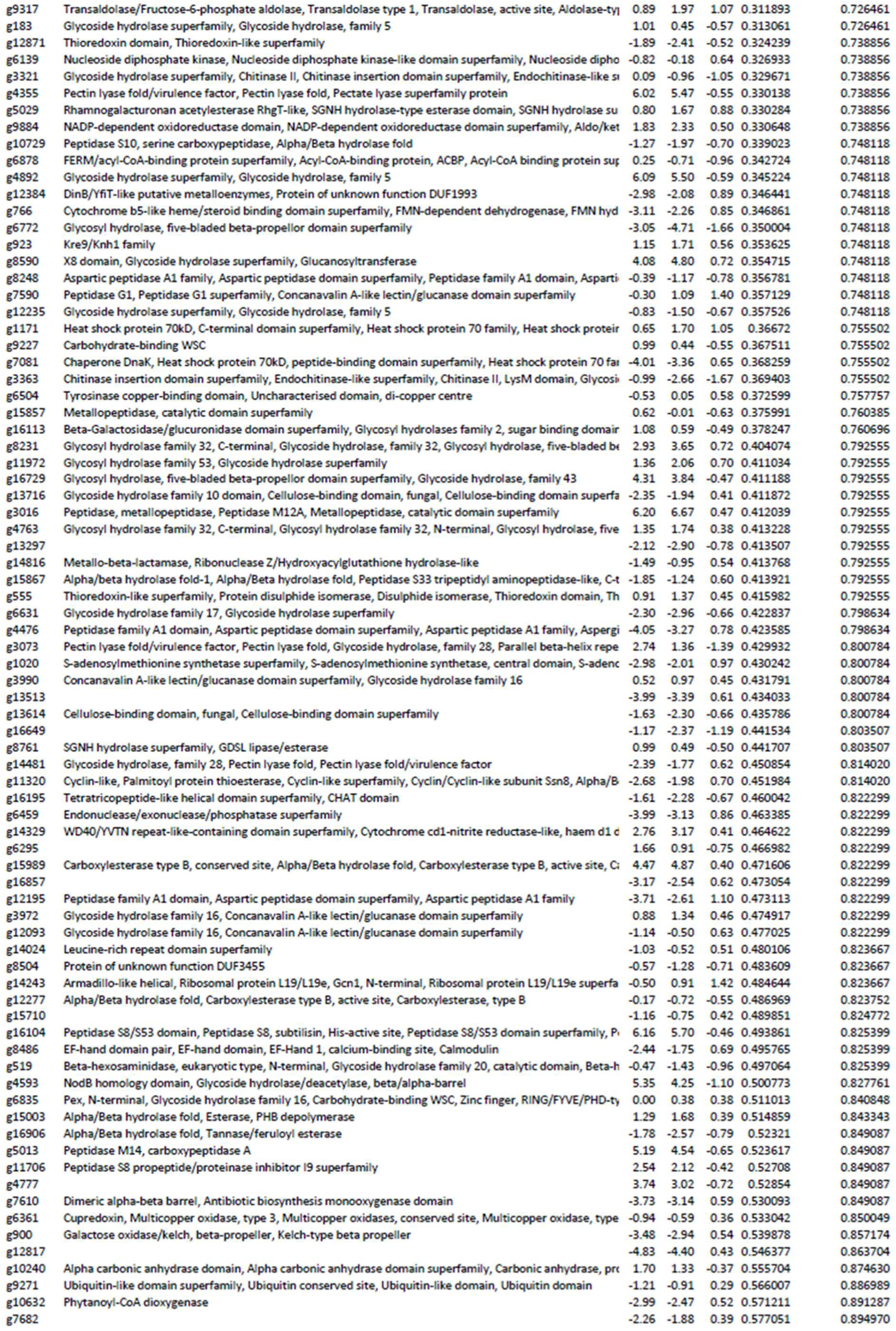
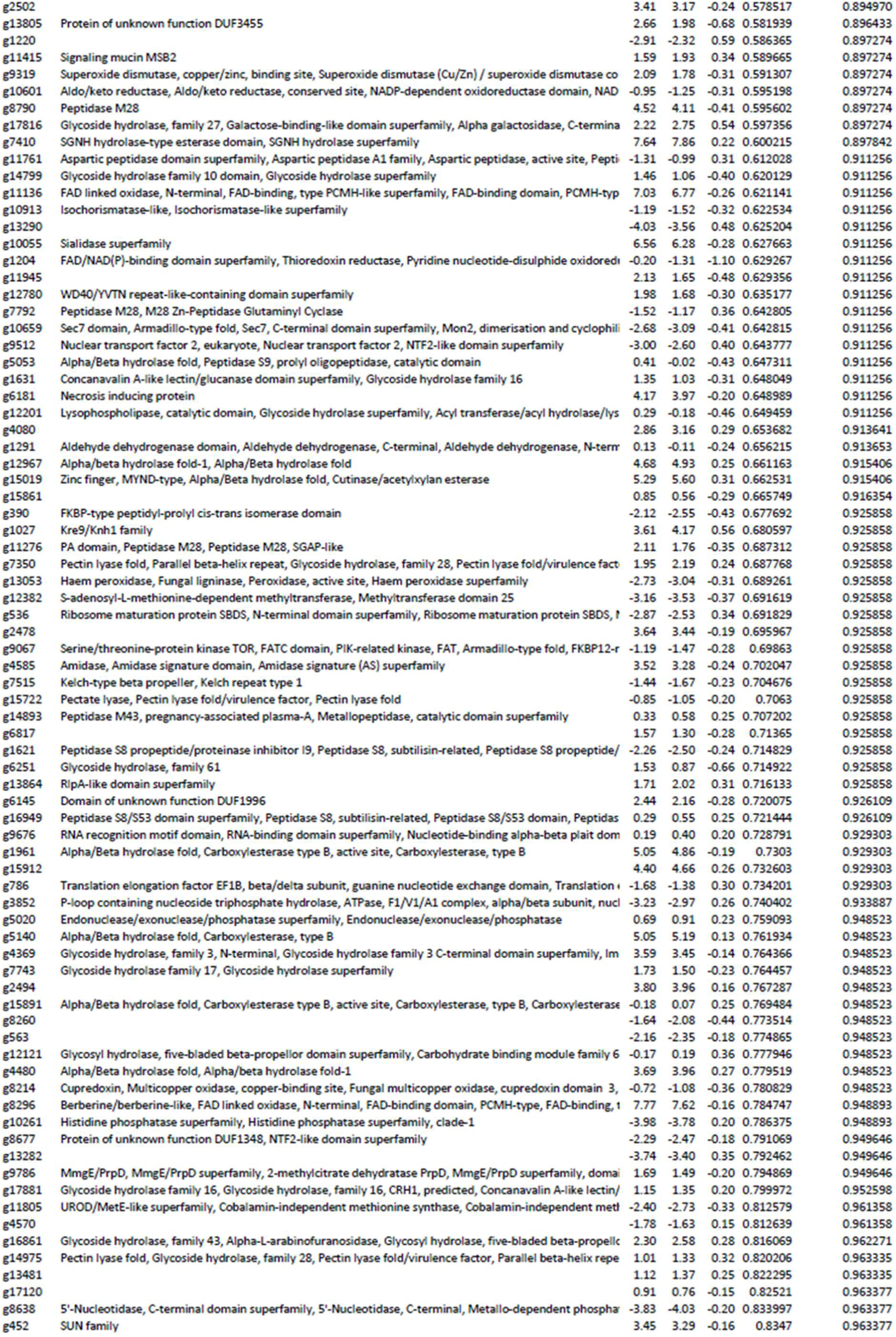
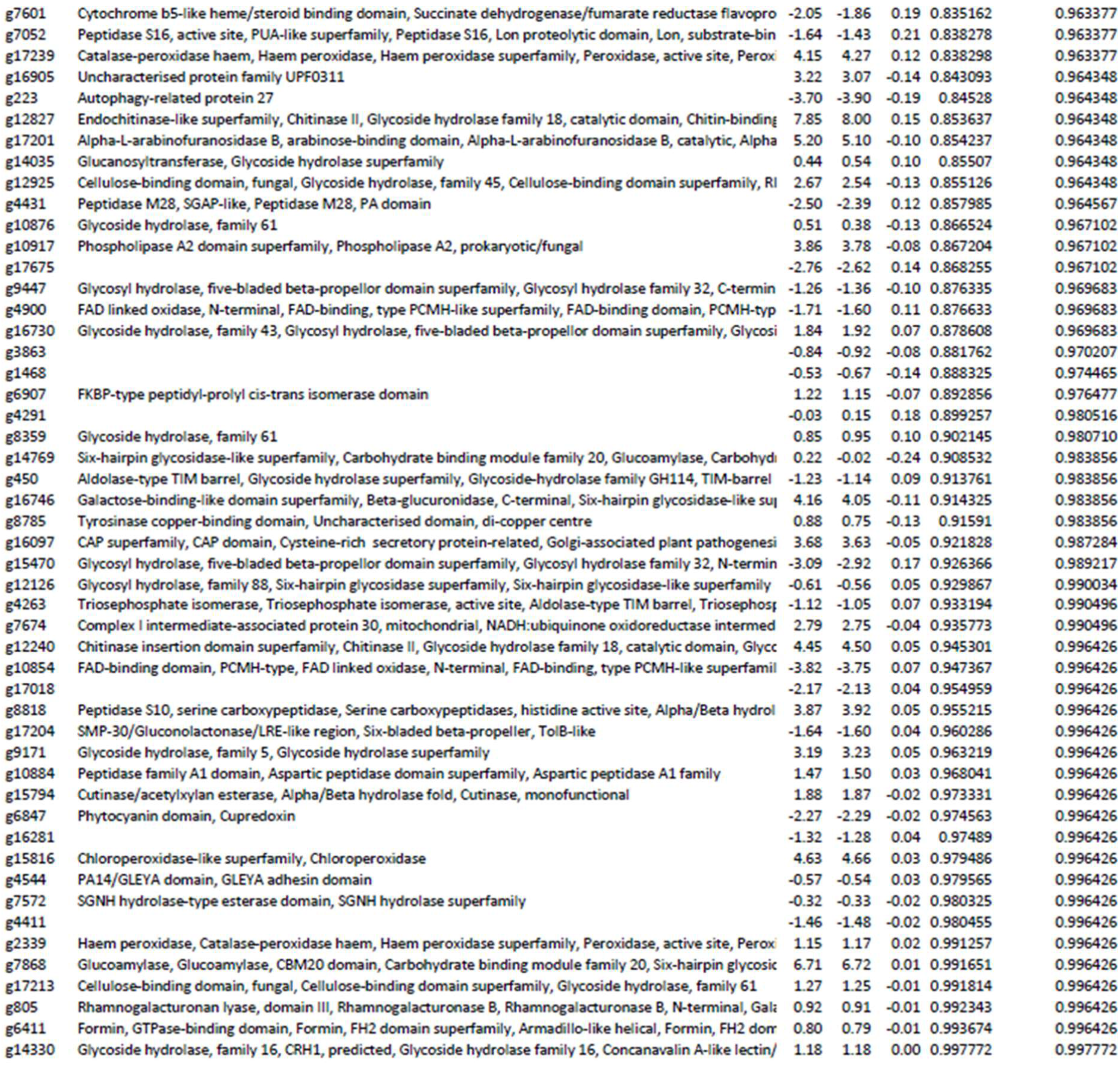
Fo5176 secretome identified proteins. Proteins identified in *clr1-1* and WT Proteins identified in the secretome of WT and *clr1-1* infected roots at 3 dpt. Proteins differentially present in *clr1-1* secretome relative to WT with a moderated T-test p<0.05 are highlighted in green and red for less or more abundant in the mutant, respectively, as in Fig 5B. For the proteins without IPR description, no homologues were found in the database to assign a putative activity.

In contrast, we found an increased presence of other virulence factors in the secretome of *clr1-1* compared to that of WT (Fig. 5B and Table 1). Interestingly, *clr1-1* releases increased amounts of a protein (g2038) annotated as a cutinase, which was reported to be essential for virulence of necrotrophic fungal pathogens (*48*). Additionally, we found an increase in pectin and pectate lyases in the *clr1-1* secretome (*49*–*51*). Similarly, a set of 8 peptidases previously described as critical virulence factors (*52*), were more abundant in the *clr1-1* secretome. These included 4 subtilases and fungalysin, among others (Table 1). The subtilases are able to degrade plant proteins with antifungal activity, such as β1-3 glucanases (*53*). The fungalysin proteins compromise host defense by cleaving class IV plant chitinases, preventing the release of the MAMP chitin and, as such, have been reported to be necessary for the virulence of several plant pathogenic filamentous ascomycetes, including Ustilago *maydis, F. oxysporum f sp. lycopersici* (Fol), *F. verticillioides* and *Colletotrichum graminicola (54–57)*. Finally, we noticed an enrichment in a chitin deacetylase, whose orthologs impede chitin-triggered immunity in cotton by converting chitin oligomers into ligand-inactive molecules like chitosan (*58*). Our data indicate that *clr1-1* reduces the plant detection of its chitin, which could explain its increased virulence and the reduced host defense (Fig 5A). To test this possibility, we performed a chitin-dependent defense priming assay as previously done for cellobiose (Fig 5b). Indeed, when roots were pre-treated with chitin, the plant susceptibility to *clr1* was indistinguishable from that observed against WT Fo5176 (Fig. 5C).

We next investigated the role of sugar availability on the up-regulation of the virulence factors enriched in the *clr1-1* secretome. We measured the expression of six of these genes (g16048, g2038,g7764,g1489,g15789 and g11385) in WT and *clr1* grown on sucrose media or on roots at 7dpt. We observed an increase in the expression in all the cases except with the subtilisin (g1489) in plants infected by *clr1-1* compared to WT, while no differences were observed between the strains grown on sucrose (Fig. 5D-E). These results suggest that *clr1-1* increases both the expression and secretion of various virulence factors to compensate for its decreased ability to utilize cellulose as a primary carbon source.

### *CLR1* is essential to complete *F. oxysporum* life cycle

Our data indicated that CLR1 is not required for Fo5176 to reach and enter the xylem and question the biological relevance of this transcription factor in Fo growth in planta. To shed light on this, we studied the temporal transcriptional regulation of *CLR1* in infected roots. *CLR1* expression increased significantly from 4dpt on, especially after 5dpt (Fig. 6A), growing exponentially until 7dpt, when Fo5176 had already proliferated in the vasculature. This suggests that cellulases might have a critical role in the final infection stages. We therefore explored CLR1 function during the final phase of the host-fungus interaction, saprophytic growth, which is an important part of the Fo life cycle. When Fo5176 was grown for 4 days on dead or living plants, we observed 3 times more expression of *CLR1* under saprophytic than under hemibiotrophic growth conditions (Fig. 6B). The last step of a fungal life cycle is the production of spores. Therefore, we tested the impact of CLR1 on microconidia production on dead seedlings. Our data showed that the WT and *clr1C* produced five times more microconidia than *clr1-1* (Fig. 6C). Taken together, these findings indicate that CLR1 is essential for saprophytic growth and spore production of Fo on dead plant tissue.

**Fig. 6.**
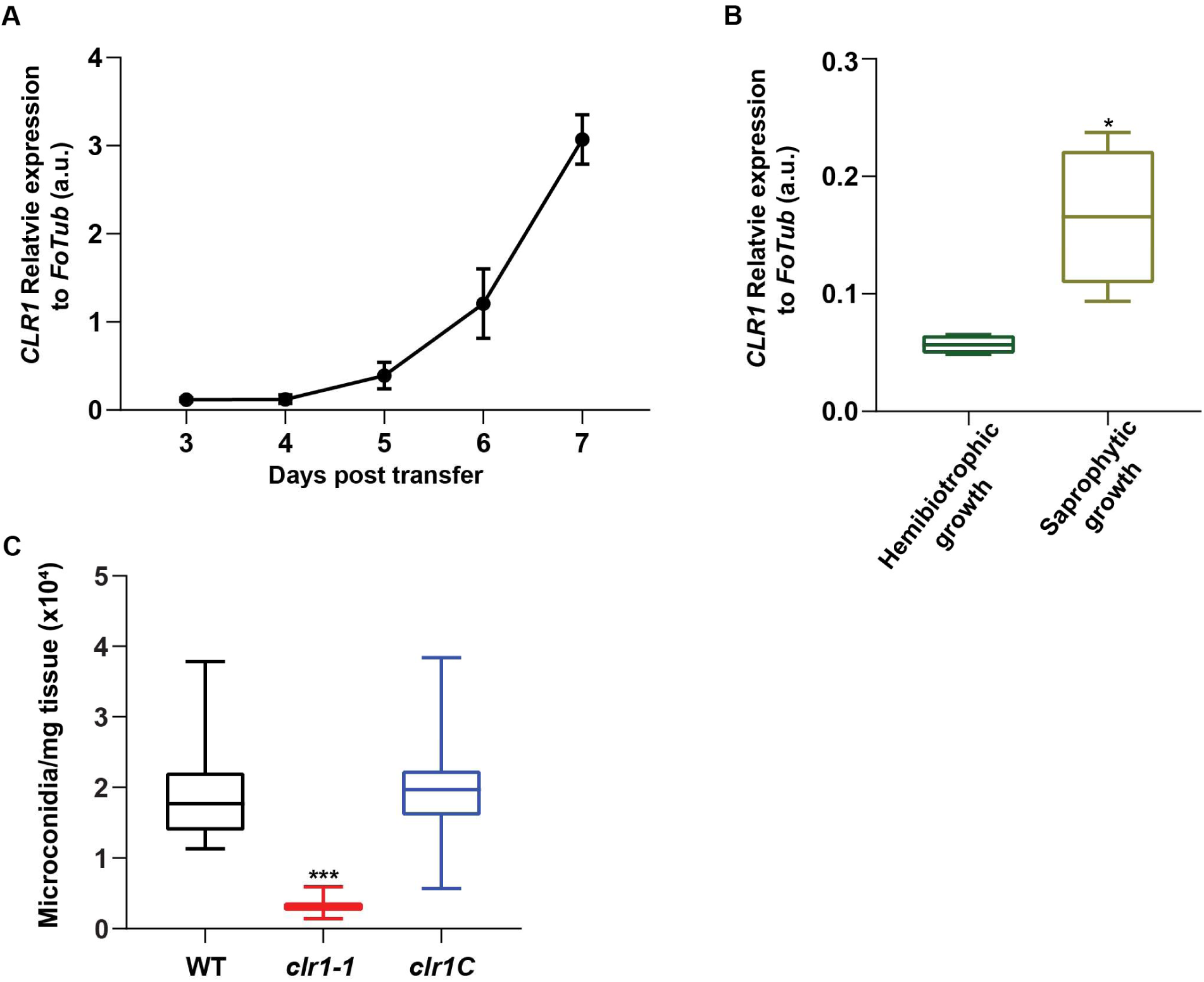
CLR1 is crucial for Fo saprophytic growth and spore production. (A) *CLR1* expression analysis relative to *FoTub* in Arabidopsis infected roots at different dpt to WT microconidia containing plates. Values represent means +/- SEM from 3 biological replicates. **(B)** *CLR1* expression relative to *FoTub* in WT or *clr1-1* grown in alive or dead plants for 4 days. Shown are box plots: center lines show the medians; box limits indicate the 25th and 75th percentiles; whiskers extend to the minimum and maximum, N≥4 biological replicates; Welch’s unpaired *t*-test; * p-value<0.05. **(C)** Microconidia production of WT, *clr1-1* and *clr1C* in dead aerial tissue of infected Arabidopsis plants. Box plots as described above. N≥10 biological replicates. Asterisks indicate statistical difference relative to WT, Welch’s unpaired t-test. ***p<0.001.

### The role of CLR1 in Fo infection is conserved among different pathosystems

The *F. oxysporum* species complex contains different host specific plant pathogens, for some of which we also found CLR1 orthologues with a protein sequence similarity of 95.69% with the Fo5165 CLR1 and the same DNA-binding sequence (Fig. 1A and 1B). Thus, we asked whether the increased virulence displayed by *clr1-1* in the Arabidopsis-Fo5176 interaction is conserved among pathosystems. To answer this question, we generated a *CLR1* null mutant in the tomato-infecting Fo, Fol4287, and the corresponding *clr1C* complemented line (Fig. S7). Similar to Fo5176, tomato plants infected with Fol strains, a higher virulence of *clr1-1* mutant compared to the WT was observed, while *clr1C* was similar to WT (Fig. 7). These results suggest that CLR1 is not essential for *F. oxysporum* infection, both in Fo5176-Arabidopsis and Fol4782-tomato interactions.

**Fig 7.**
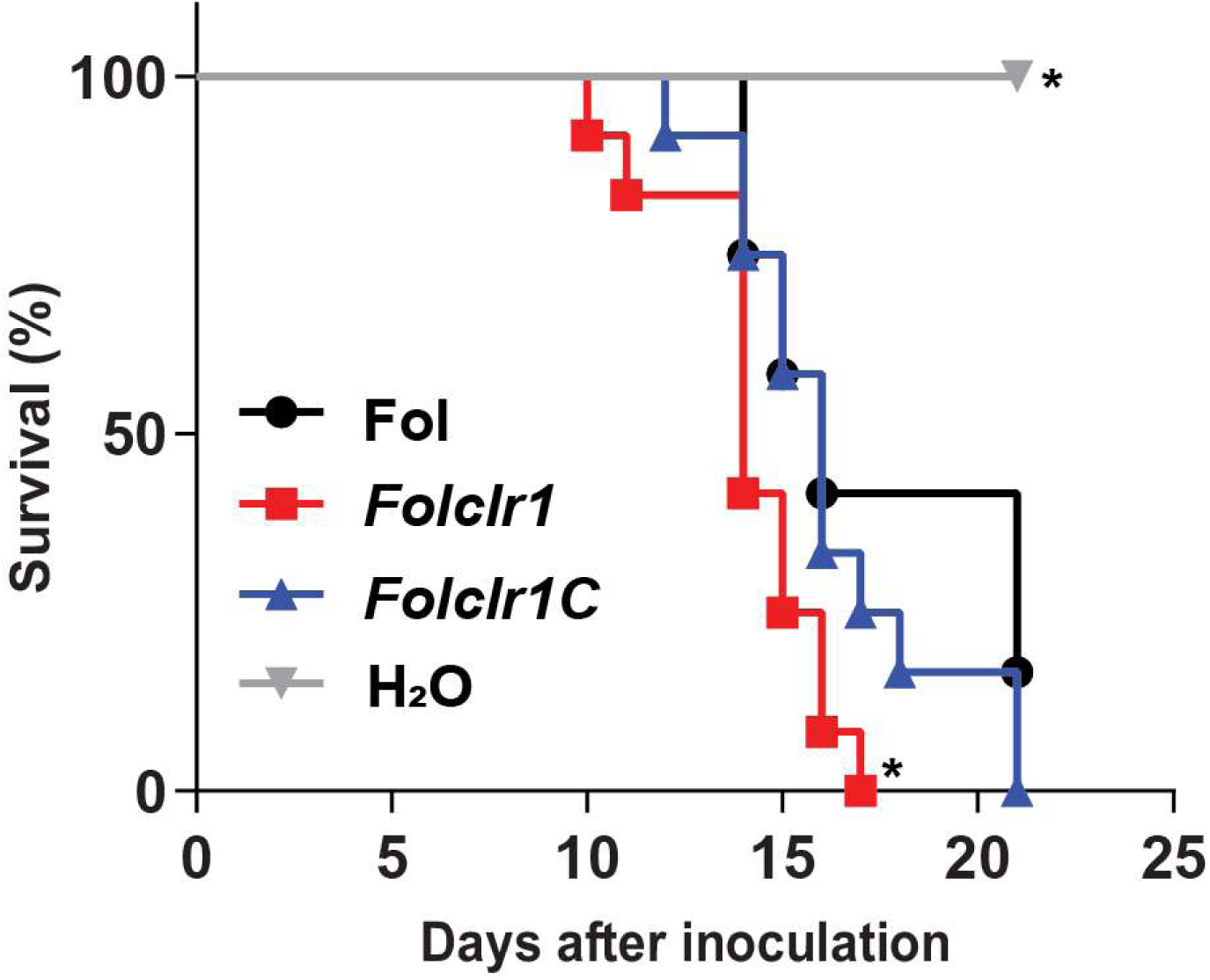
The lack of CLR1 also increases Fol4287 pathogenicity in tomato plants. Kaplan–Meier plot showing the survival of tomato plants grown in vermiculite and dip-inoculated or not (H_2_O) with *Fol, Folclr1* or *Folclr1C* microconidia. N=15 plants from one independent experiment. \**P* < 0.05 versus *Fol* alone according to log-rank test. The experiment was repeated 3 times with similar results.

**Fig. 8.**
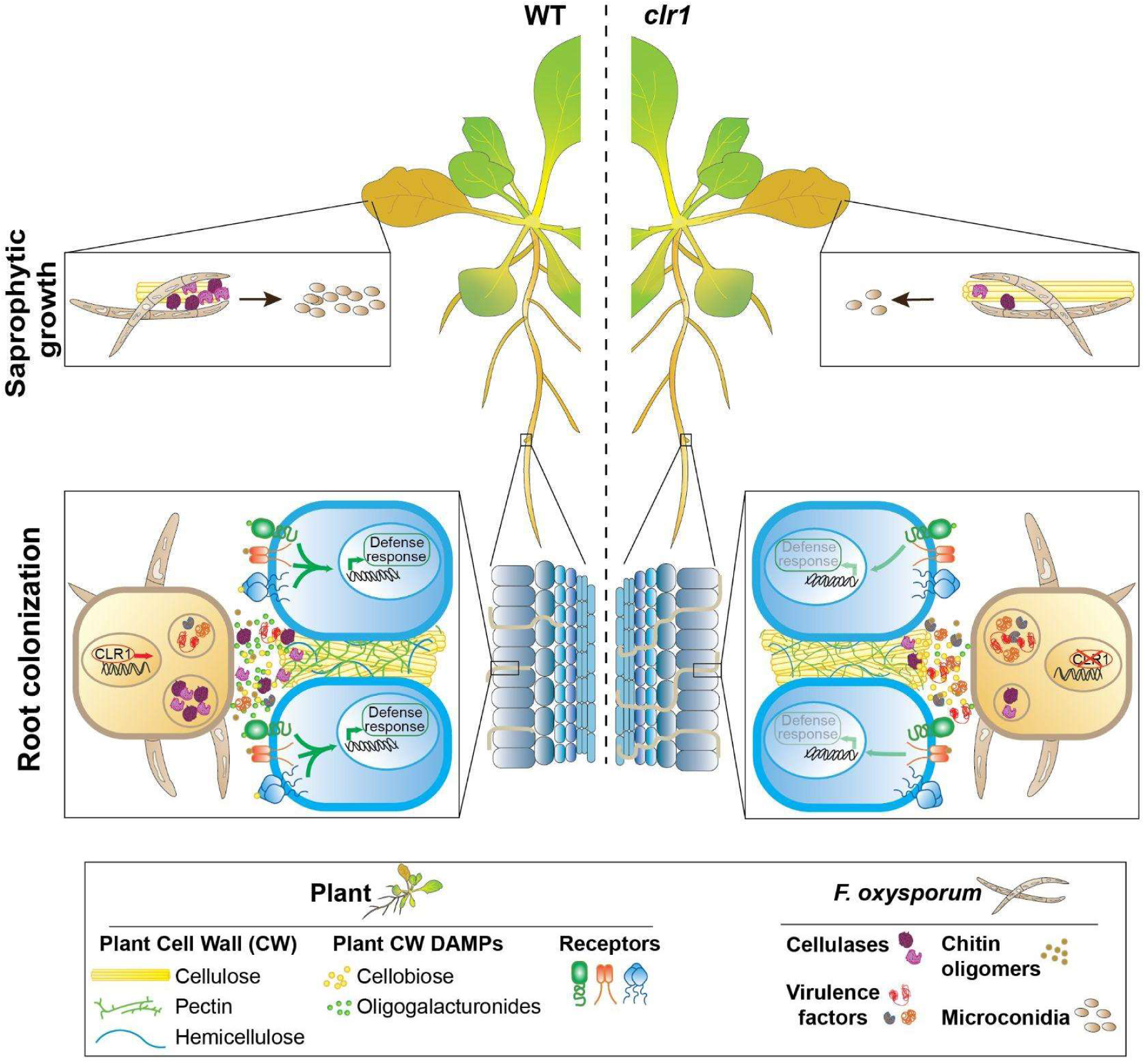
Model of CLR1 role in the life cycle of *F. oxysporum* in planta. Bottom part: During root apoplast colonization, CLR1 is required to degrade and consume the plant cellulose. In its absence, the *clr1* mutant secretes more virulence factors and produces less oligogalacturonides, which are pectin-derived damage-associated molecular patterns (DAMPs). The virulence factors secreted by *clr1* also reduce the production of chitin-monomers perceived as microbe-associated molecular patterns by the plant. Consequently, the host recognizes *clr1* less efficiently than WT and *clr1* can colonize faster the xylem. Green arrows represent signalling cascade activation, their intensity reflects how much it is triggered. Upper part: Once the plant is dead, Fo needs to grow saprophytically and needs CLR1 to use cellulose as carbon source and produce microconidia.

## Discussion

Microbes need to loosen and digest the plant CWs in order to colonize their hosts. As the major and most recalcitrant CW component, cellulose is a key target for a plethora of transcriptionally coordinated microbial CWMPs (*34, 36, 37, 59*–*63*). In this work, we studied the function of cellulases in the life cycle of a plant fungal pathogen. The identification and characterization of CLR1 in Fo confirms the conservation of this transcription factor as a master regulator of cellulose degradation in ascomycetes (Fig. 1, fig. 2H and fig. S3) (*8, 36, 37, 43*). Our work expands the current knowledge on the role of CLR1 during plant biomass degradation by demonstrating its function *in planta*. Interestingly, genes described in *N. crassa* as CLR1-dependent in cultures with cellulose as the only carbon source behaved similarly in Fo during root infection, suggesting that both metabolic situations require similar levels of CWMPs (Fig 2G; (*37*)). Unexpectedly, loss of CLR1 led to a significant increase in virulence both in the Arabidopsis and tomato pathogenic forms of Fo (Fig 2, fig. S4, and fig. 7). Although the *clr1* mutant is not completely impaired in cellulases secretion (Table 1), our data indicate that Fo not only does not need their CLR1-dependent cellulases, but benefit from their absence to infect their hosts. More precisely, we show here that *clr1* crosses the root epidermal layer and reaches the vascular system of Arabidopsis roots faster, while degrading less crystalline cellulose, than WT (Fig. 2A-F, and H). This indicates that crystalline cellulose degradation is not a requirement for Fo to grow through the apoplast of the root layers and that the impairment of *clr1* in cellulose degradation is a consequence of the notable decrease in the expression and secretion of most, but not all cellulolytic enzymes (Fig. 2G, fig. 5B and 5E, and Table 1).

Our previously reported transcriptomic data indicated that the days 2 and 3 of the root-Fo5176 interaction are critical time points for disease development (*34*). The drastic decrease of sucrose levels detected in infected roots at these time points (Fig 4A-C) suggests that the initial steps of root colonization have an extraordinary impact on plant metabolism. Accordingly, we observed a significant impairment in plant response to *clr1* infection at these same time points (Fig 3A), indicating that the lack of a canonical set of CWDEs stunts the perception of the pathogen by the plant. However, although pre-treatment with cellobiose reduced the virulence of the *clr1* mutant (Fig3B), its detection in higher amounts in *clr1*-infected roots as compared to WT excludes this cellulose degradation product as the main reason for the delay in the perception of *clr1* by the plant. The high amounts of cellobiose detected in *clr1*-infected roots and the reduced capacity of *clr1* to grow on cellobiose as the only carbon source could be explained by the markedly reduced expression levels of the enzyme β-glucosidase, which is required for the final step of cellobiose hydrolysis (*64*). Importantly, the identification of cellobiose as a CW degradation product during microbial colonization confirms its predicted but never confirmed function as DAMP (*27*). Together with cellobiose, we identified various OGs in Fo-infected roots (Fig. 4B-D). Interestingly, most of the OGs identified in roots infected either with WT and *clr1* were acetylated, which differs from what was reported in leaves infected by the necrotrophic fungus *B. cinerea (65)*. Aerial and soil-borne plant pathogens might thus use a different battery of CWDEs to adapt to the different CWs encountered in leaves and roots, with some functional similarities since certain OGs such as GalAox were detected in both infected organs. Acetylated OGs have been shown to impair the activity of some pectinases, eliciting the plant defense response (*66, 67*). Thus, the reduced amount of GalAc_2_ox and Gal_4_Ac detected in *clr1*-infected roots (Fig. 4D and E) might explain the delay in fungal perception by the plant.

Plant pathogens modulate their secretome to adapt to the host and the available nutrient sources (*8, 68*). Together with the decreased amount of cellulases in the *clr1* secretome, we found an enrichment of secreted virulence factors, which might partially explain the fact that co-infection with WT does not reduce the hypervirulence of the mutant (Fig. S6A and B). Intriguingly, the most enriched protein in the *clr1* secretome was a putative cutinase. Cutinases are essential for virulence of aerial plant pathogens (*48*) where the cuticle is a substantial barrier on the leaf and stem surfaces. In roots, only the root caps have been shown to have a cuticle (*69*). However, both pathogenic fungi and mutualistic mycorrhizae recruit the fatty acid biosynthesis program to facilitate host invasion. As reported for these microbes (*34, 70*), we observed an upregulation of the expression of these fatty acid biosynthesis genes in Fo during Arabidopsis root infection. In addition, cutinases from plant pathogens were shown to cleave suberin *in vitro* (*71*), a plant compound that prevents the spread of microbial pathogens (*72*). Because suberin blocks apoplastic transport both at the endodermis and at the root surface (*73*), the increased amounts of cutinase might explain the ability of *clr1 to* cross the root epidermal layer and reach the xylem faster than WT (Fig 2).

We also found a set of peptidases enriched in the *clr1* secretome, some of which have been reported to affect host responses by degrading defense related proteins (*52, 74*). Among them, fungalysins have been reported to be involved in cleavage of plant chitinases and therefore be essential for the pathogenicity of different fungi, including Fol *(56, 75)*. Interestingly, the *clr1* mutant also seems to undermine host defense responses by secreting more chitin deacetylases, which should decrease the amount of MAMPs available for plant perception (*58*). In fact, we found that pretreatment with chitin oligomers reduces *clr1* virulence to WT levels, implying that not only DAMP, but also MAMP perception is severely compromised during *clr1* infection. The other group of virulence factors significantly enriched in the *clr1* secretome are the pectin and pectate lyases, which have a fundamental role in vascular wilt diseases (*50*). These enzymes generate unsaturated OGs that might be used efficiently as nutrients by Fo because most OGs detected in the secretomes were saturated. In line with this, the only unsaturated OG identified in the assay, GalA_3_Ac-H_2_Oox, was less abundant in the *clr1* secretome (Fig. S5). Interestingly, we did not find differences in the levels of polygalacturases (PGs) despite the higher levels of saturated OGs observed in infected roots. Most likely, the higher levels of pectin and pectate lyases detected might provide additional substrates for PGs, whose activity would not be limitant in this reaction. Indeed, the overaccumulation of subtilisin in the *clr1* secretome could create more cleavage sites for the PGs by activating type I pectin methyl esterases (*76*). The lower amount of saturated OGs observed in *clr1*-infected compared to WT-infected roots might thus be a consequence of (a) the higher cellulose degradation during WT infection that makes the cellulose-embedded pectin more accessible for deconstruction by pectinases or (b) the increased capacity of *clr1* to uptake, and use these OGs. We favour the second hypothesis considering that the majority of pectins is not bound to cellulose (*77*). Overall, the increase in the secretion of different virulence factors might explain the delay in the perception of *clr1* by the plant at the early stages of infection (Fig. 3Aa).

Sucrose was highly available for Fo at the beginning of its interaction with the root (2dpt), when Fo is entering the epidermal apoplast (*34*), while the CW-derived oligomers were not detectable (Fig. 4). At 3dpt when some hyphae have reached the cortical apoplast (Fig.2F (*34*)), sucrose levels had decreased dramatically and cellobiose and OGs were detectable, indicating that from 2 to 3 dpt the CW becomes the primary carbon source for the fungus. At this stage, energy acquisition might become limiting in *clr1* due to its reduced capacity to degrade cellulose. In this metabolic context, the enrichment in pectin and pectate lyases detected in the *clr1* secretome might represent a metabolic adaptation to exploit energy through the alternative carbon source pectin. Moreover, the higher levels of peptidases secreted by *clr1* could also provide additional carbon and nitrogen sources to the fungus (*78*). This hypothesis is supported by the fact that the increase in the expression of virulence factors in *clr1* compared to WT was only observed during growth *in planta* but not on carbon-rich axenic medium (Fig. 5D and E).

Finally, the composition of the medium strongly affected conidia production by Fo. A positive role of cellulose in conidia formation was reported previously (*79*), accordingly, *CLR1* expression was upregulated starting at 4dpt, when Fo has already entered the xylem (Fig. 6A and 2A), and remained significantly high when the fungus was growing on dead plants compared to alive ones (Fig. 6B). Thus, CLR1 activation is likely a way to prepare the fungal metabolism for the next step in the Fo life cycle, i.e. formation of conidia on the decaying plant tissue which will spread the fungus to encounter new hosts. Indeed, we showed that *clr1* was severely compromised in asexual reproduction because microconidia formation on dead plants decreased drastically (Fig. 6B and 6C). Our data suggest that cellulose degradation is fundamental for the pathogen to fully exploit host resources for the production of dissemination structures.

In conclusion, we show for the first time that the impairment in cellulose degradation does not pose any obstacles for plant infection by the vascular pathogen Fo. We further demonstrate that this organism exhibits an extraordinarily high degree of metabolic plasticity that compensates the decrease in cellulase production during host colonization by triggering the expression of alternative CWDEs. However, the deficiency in obtaining energy from cellulose significantly decreases the capacity of the fungus to replicate, disseminate, and invade new hosts. Our findings imply that CLR1 is highly conserved among ascomycetes because it is vital for saprophytic survival in natural environments. Evolutionary pressure thus makes the use of cellulases imperative for successful completion of the pathogen’s life cycle.

## Material and Methods

### Plant material and growth conditions

Arabidopsis thaliana ecotype Col-0 and tomato Monika (provided by Syngenta Seeds, Almeria, Spain) were used in the analysis of fungal pathogenicity. Growth conditions were 16-h light (24°)/8-h dark cycle at 21º C for all Arabidopsis experiments. In the case of plate experiments, seedlings were grown upright with ½MS media (pH adjusted to 5.7 with KOH but without buffering). When the experiment was performed in a hydroponic system, the seeds were germinated on 2 mm foam plugs floating in 330 ml pots on ½MS + 1% sucrose media at pH 5.7 adjusted by KOH. The media was exchanged 6 days after germination to ½MS and seedlings were further grown. Tomato cultivar susceptible cultivar Monika (provided by Syngenta Seeds, Almeria, Spain) planted in vermiculite, and maintained in a growth chamber (28 °C; photoperiod 14 h light/10 h dark)

### Fungal strains and culture conditions

*Fusarium oxysporum* strain 5176 (Fo5176) was originally isolated in Australia from infected Arabidopsis plants (*38*). The strains were routinely cultured in potato dextrose broth (PDB) at 28°C with orbital shaking at 170 rpm. Where necessary the following antibiotics were added to the culture medium: hygromycin B (55 μg/ml), G418 (100 μg/ml) and phleomycin (5.5μg /ml). For microconidia collection, 3 to 5 day-old cultures were collected by filtration through a nylon filter (Monodur; mesh size 10 μm). Filtrates were centrifuged at 12000 g for 10 min, the pellet containing the microconidia was washed using deionized water, resuspended in water to reach the desired concentration.

### Identification of CLR1 in Fusarium species

Fusarium genes encoding the predicted CLR1 proteins were identified by sequence similarity searches against the *Neurospora crassa* CLR1 (NCU07705) (*36*) using BLASTp (http://blast.ncbi.nlm.nih.gov/). The software CLC Genomic Workbench v.12 was used to perform the comparison analysis of CLR1 sequence in different ascomycetes.

### Fungal transformation

As described above, we identified the CLR1 proteins in Fo5176 (FOXB_08021) and Fol4287 (FOXG_08626). Targeted gene deletion of the entire *CLR1* gene in the *Fo5176-pSIX1::GFP* background (*33*) and the *Fol4287*-3X*mClover3* genetic backgrounds was performed using the split marker method (*80*) with the neomycin resistance cassettes following the protocol previously described (*63*). For the complementation of the *clr1* deletion mutants, a co transformation with the native *CLR1* in the phleomycin resistance cassette was performed as reported (*81*). The oligonucleotides used to generate PCR fragments for gene replacement, complementation or identification of mutants are listed in Table 1. PCRs were routinely performed with the High-Fidelity Template PCR system (Roche Diagnostics, Barcelona, Spain) using an MJ Mini personal thermal cycler (Bio Rad, Alcobendas, Spain). The amplified flanking sequences were PCR fused with partially overlapping truncated versions of the neomycin (Neo^r^). Transformants were purified by monoconidial isolation (*82*).

### Fungal Growth in different carbon sources

Fo5176 was grown on ½MS media (Murashige and Skoog media, Difco). Carbon sources were added to 0.5% wt/vol. Conidia were inoculated into 3 ml liquid media at 10^6^ conidia/mL and grown at 28º in dark and shaking (180 rpm), 3 days for sucrose and cellobiose (Fluka) and 7 days for cellulose (Sigmacell, cellulose type 50). After 1 day drying at 60º, the material was weighted or its nucleic acids were extracted. For the nucleic acid extraction, we followed a previous protocol (*83*). The nucleic acids were quantified by spectrophotometry using a NanoDrop.

### Fungal growth in *in vitro* stress conditions

Drops containing serial dilution (1×10^7^, 1×10^6^ and 1×10^5^) of freshly obtained microconidia were spotted on YPD agar plates supplemented with Congo red (5 μg/mL) or 5 μg/mL CalcoFluor White prepared in 0.5% KOH for testing cell wall stress as in (*84*). Minimal media was used for evaluating growth under nutrient deficiency, for (*82*)1M sorbitol was used to test hyperosmotic stress. Plates were incubated at 28 °C for 2 days in general or 4 days for salt stress and Calcofluour white before imaging..

### Arabidopsis plant infection assays

Arabidopsis infection assays were performed as described previously (*33, 41*). In summary, 8 days-old seedlings grown as described above were transplanted to plates with 100 μl of 10^7^ microconidia/ml of Fo5176-pSIX1. The number of vascular penetrations per root was analyzed using the signal from the GFP under a stereomicroscope, when it showed a clear, linear and root central pattern. The experiment in which the plants were pretreated with cellobiose 100 μM or chitin 100μg/ml were based on (*27*). Roots of 10 days old seedlings were treated with cellobiose or chitin for 25 minutes, hereafter they were transferred on a ½MS media plate with 3 days-old Fo mycelia. The number of vascular penetrations were monitored as explained above.

### Tomato plant infection assays

Tomato root infection with *Fol* was performed as previously described by (*82*). Briefly, 2 week-old seedlings were inoculated with *Fol* by immersing the roots into a fungal microconidia suspension (5×10^5^ conidia/ml), planted in vermiculite, and maintained in a growth chamber (28 °C; photoperiod 14 h light/10 h dark). Plant survival was recorded daily up to 29 days, as previously described (*85*), calculated by the Kaplan–Meier method, and compared among groups using the log-rank test. All infection assays included 10 plants per treatment.

### Confocal microscopy

Plates infection assays were performed as described above, at 3 dpt images were taken with a Zeiss LSM Axioobserver microscope, using the LD C-Apochromat 40x / 1.1 W Korr M27 objective and Immersol W (Zeiss) between lens and coverslip. GFP (fungus) was excited at 488 nm and emitted fluorescence was detected at 514 nm. The roots were stained with propidium iodide (PI; 10 μM, Sigma-Aldrich) and imaged with 536 nm and 617 nm excitation and emission wavelength, respectively.

### Cellulose, OGs and hexoses and quantification in infected roots

Plates infection assays were performed as indicated above. For cellulose quantification, roots were harvested at 7dpt and processed as described before to measure the crystalline cellulose (*33, 86*). To quantify OGs and hexoses, roots collected at different dpt were weighted, covered with 100% ethanol for several days and then dried using a freeze-dryer (Christ, Alpha 2-4). The roots were grinded in water (1ml), centrifuged (10 minutes at 10.000g) and the supernatant was collected to be analyzed via High-performance size-exclusion chromatography (HP-SEC-MS). The analysis was performed as indicated before (*65*). Briefly, samples were diluted at 1 mg/ml in ammonium formate 50 mM, formic acid 0.1%. Chromatographic separation was performed on an ACQUITY UPLC Protein BEH SEC Column (125Å, 1.7 μm, 4.6 mm X 300 mm, Waters Corporation, Milford, MA, USA). Elution was performed in 50 mM ammonium formate, formic acid 0.1% at a flow rate of 400 l/min and a column oven temperature of 40 °C. The injection volume was set to 10 l. MS-detection was performed in negative mode with the end plate offset set voltage to 500 V, capillary voltage to 4000 V, Nebulizer 40 psi, dry gas 8 l/min and dry temperature 180 °C. Major peaks were annotated following accurate mass annotation, isotopic pattern and MS/MS analysis, as previously described (*45*).

### Real-time quantitative PCR

LN2 frozen samples were grinded using a TissueLyser II (Quiagen) and glass beads. 1 μg of total RNA/sample extracted by Isol-RNA lysis reagent (5 PRIME) was used to generate first-strand cDNA using Thermo Scientific Maxima™ H Minus cDNA Synthesis Master Mix with dsDNase (Thermofisher), following manufacturer’s instructions. Quantitative PCR reactions were performed in a LightCycler 480II apparatus (Roche) using Fast SYBR Green Master Mix (Thermofisher) in a 10 μl reaction. Relative transcript levels were quantified with respect to a reference gene *GAPDH600B* for plants (*87*) (Czechowski et al. 2005) or *FoβTUB* (F*OXG_06228*) for Fo5176 (*33*). The 2^ΔCT^ method was used to quantify the relative expression of each gene. Primers are indicated in Table S1.

### Protein identification in Fo5176 secretome

10-day old hydroponically-grown seedlings were treated with 20 μl of 10^7^ microconidia/ml of either Fo5176 WT or *clr1*, as described before (*34*). At 3dpt,, the roots were removed and the liquid media was collected. We next followed the protocol from (*88*) with some modifications: to remove big particles present in the media, we filtered it with a 45 microns sterile filter (Starlab) before concentration with. centricons (Amicon Ultra 3k, Merck Millipore) to 1.5 ml volume. 150 microliters of these 1.5 ml were sent to proteomic analysis.

For each sample, proteins were precipitated with trichloroacetic acid (TCA; Sigma-Aldrich) at a final concentration of 5% and washed twice with cold acetone. The dry pellets were dissolved in a 45 μl buffer (10 mM Tris + 2 mM CaCl2, pH 8.2). Reduction and alkylation of the proteins was performed by adding 2 mM of Tris(2-carboxyethyl) phosphin –hydrochlorid (TCEP) and 15 mM of iodoacetamine (IAA). After 30 min at 60°C the samples were cooled to room temperature and 4 μg of Sequencing Grade Trypsin (Promega) for digestion were added. The digestion was carried out at 37°C for 4 hours. The samples were dried to completeness and re-solubilized in 20 μl of 3% acetonitrile, 0.1% formic acid for LC-MS/MS analysis. Before injection the samples were diluted 1:20 in the same solvent.

Mass spectrometry analysis was performed on an Orbitrap Fusion Lumos (Thermo Scientific) equipped with a Digital PicoView source (New Objective) and coupled to a M-Class UPLC (Waters). Solvent composition at the two channels was 0.1% formic acid for channel A and 0.1% formic acid, 99.9% acetonitrile for channel B. For each sample 1 μL of diluted peptides were loaded on a commercial MZ Symmetry C18 Trap Column (100Å, 5 μm, 180 μm x 20 mm, Waters) followed by nanoEase MZ C18 HSS T3 Column (100Å, 1.8 μm, 75 μm x 250 mm, Waters). The peptides were eluted at a flow rate of 300 nL/min by a gradient from 5 to 22% B in 80 min, 32% B in 10 min and 95% B for 10 min. Samples were acquired in a randomized order. The mass spectrometer was operated in data-dependent mode (DDA) acquiring a full-scan MS spectra (300−1’500 m/z) at a resolution of 120’000 at 200 m/z after accumulation to a target value of 500’000. Data-dependent MS/MS were recorded in the linear ion trap using quadrupole isolation with a window of 0.8 Da and HCD fragmentation with 35% fragmentation energy. The ion trap was operated in rapid scan mode with a target value of 10’000 and a maximum injection time of 50 ms. Only precursors with intensity above 5’000 were selected for MS/MS and the maximum cycle time was set to 3 s. Charge state screening was enabled. Singly, unassigned, and charge states higher than seven were rejected. Precursor masses previously selected for MS/MS measurement were excluded from further selection for 20 s, and the exclusion window was set at 10 ppm. The samples were acquired using internal lock mass calibration on m/z 371.1012 and 445.1200. The mass spectrometry proteomics data were handled using the local laboratory information management system (LIMS) (*89*).

For protein identification and label free protein quantification, the acquired raw MS data were processed by MaxQuant (version 1.6.2.3), followed by protein identification using the integrated Andromeda search engine (*90*). Spectra were searched against a provided Fo5176 database (*40*) concatenated to the Araport database (https://www.arabidopsis.org/download/index-auto.jsp?dir=%2Fdownload_files%2FSequences%2FAraport11_blastsets, version 2020-06-18), concatenated to its reversed decoyed fasta database and common protein contaminants. Carbamidomethylation of cysteine was set as fixed modification, while methionine oxidation and N-terminal protein acetylation were set as variables. Enzyme specificity was set to trypsin/P allowing a minimal peptide length of 7 amino acids and a maximum of two missed-cleavages. MaxQuant Orbitrap default search settings were used. The maximum false discovery rate (FDR) was set to 0.01 for peptides and 0.05 for proteins. Label free quantification was enabled and a 2 minutes window for match between runs was applied. In the MaxQuant experimental design template, each file is kept separate in the experimental design to obtain individual quantitative values. Protein fold changes were computed based on Intensity values reported in the proteinGroups.txt file. A set of functions implemented in the R package SRMService (*91*) was used to filter for proteins with 2 or more peptides allowing for a maximum of 4 missing values, and to normalize the data with a modified robust z-score transformation and to compute p-values using the t-test with pooled variance. If all measurements of a protein are missing in one of the conditions, a pseudo fold change was computed replacing the missing group average by the mean of 10% smallest protein intensities in that condition.

### Fo5176 microconidia production in Arabidopsis

We performed plate infection assays as indicated above. 5 days after the plant was dead, we weighed the whole plant and we added 1.2 ml of H_2_O. The samples were shaken gently for 1 minute and the plant material removed. The number of microconidia in solution counted with the Thoma cell counting chamber was shown per mg of dead material.

### Statistical analysis

Statistical analyses were performed and data plotted using GraphPad Prism 9.0.0 (GraphPad Software, Inc). Each figure legend indicates the statistical analysis which was performed and the level of significance.

## Acknowledgements

We thank all members of the Plant Cell Biology laboratory at ETHZ, especially Apolonio Huerta, Christopher Kesten, Oliver Terret, and Alexandra Menna, for technical support and fruitful scientific discussion. We are also very grateful to Niko Geldner (UNIL) for fruitful scientific discussions and to Paolo Nanni and Laura Kunz (Functional Genomics Center Zürich, FGCZ) for technical support during the generation and analysis of the proteomic data. We thank the ScopeM (ETH Zurich) for their service in the confocal microscopy.

## Author contributions

Conceptualization: CSR

Methodology: FMGA, SV, AV, SD, JCM

Investigation: FMGA, SV, AV, SD, SM, GSA, JCM

Visualization: FMGA, SV, AV, SD, GSA,

JCM Supervision: CSR and ADP

Writing—original draft: FMGA and CSR

Writing—review & editing: FMGA, SV, AV, SD, JCM, ADP and CSR

## Data and materials availability

All data are available in the main text or the supplementary materials.

## Figures and Tables

**Fig. S1.**
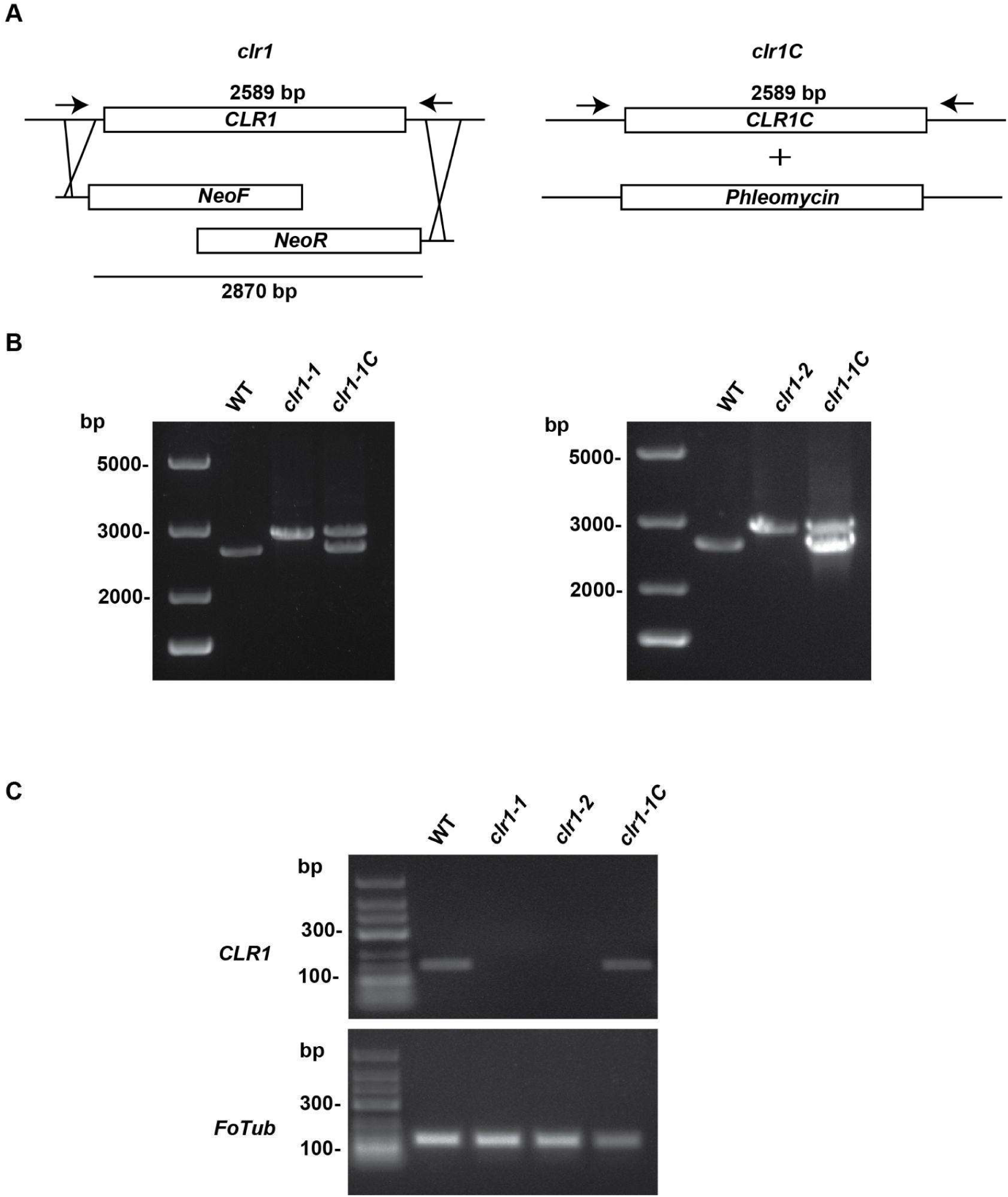
Identification of Fo5176 *clr1* mutants and *clr1C* complemented. **(A)** Left panel, schematic representation of targeted *CLR1* gene replacement with the neomycin cassette (Neo^r^) using the split-marker method. Right panel, complementation of the *clr1* mutants by co-transformation with the phleomycin resistance cassette (Phleo^r^) and the WT *CLR1* gene. **(B)** PCR screening to identify *clr1* and *clr1C* mutant strains. Genomic DNA of the indicated fungal strains was amplified by using the primer pair ForCLR1genotyping and RevCLR1genotyping (indicated with arrows in panel A). The expected PCR fragments for *clr1 or clr1C* mutants are 2589 or 2870 bp, respectively. **(C)** *CLR1* expression measured by Rt-PCR in the indicated strains using the primers pair CLR1rtpcrfor and CLR1rtpcrrev. *FoTub* expression was used as reference.

**Fig. S2.**
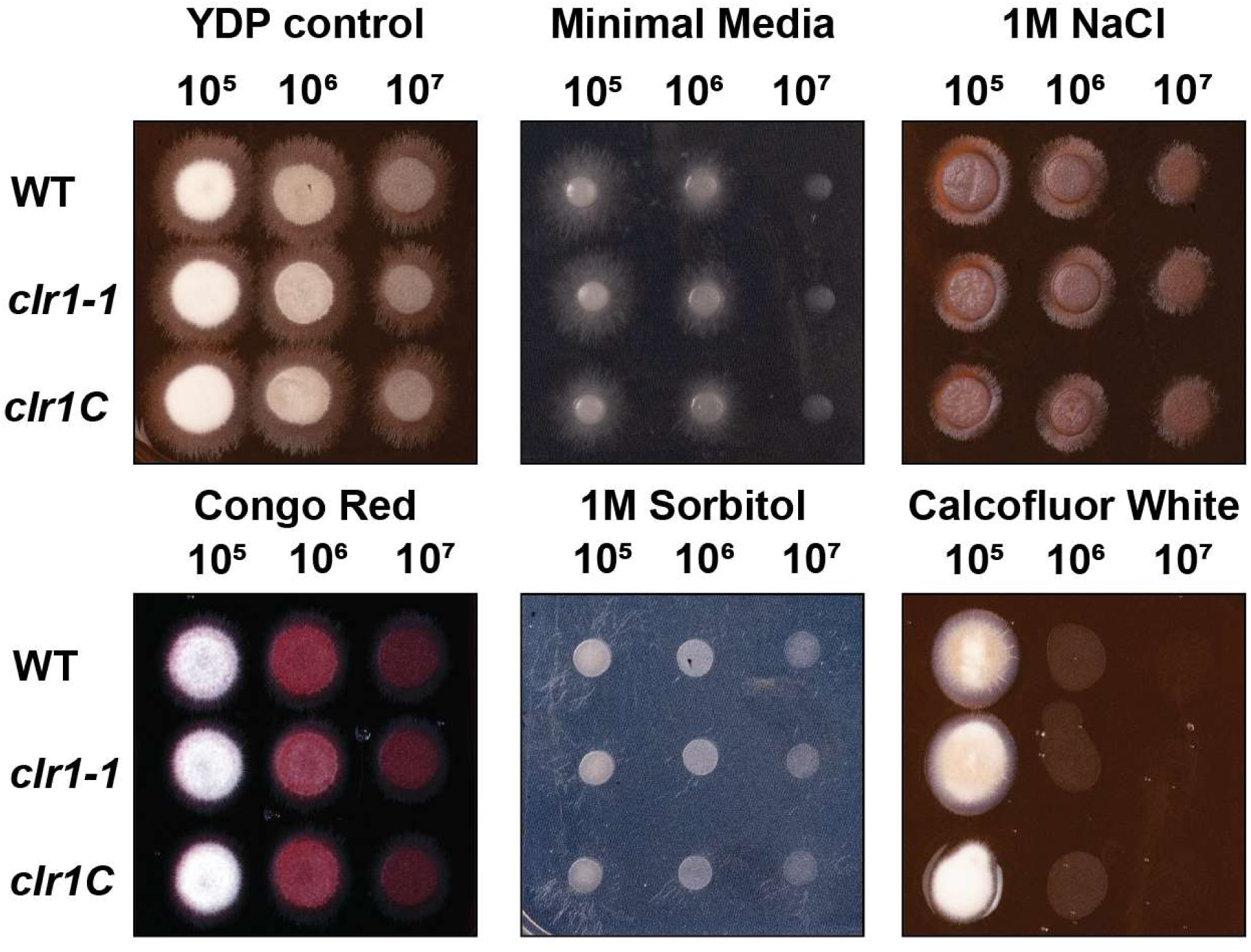
*CLR1* deletion does not alter Fo growth on different stress media. Representative images of WT, *clr1-1*, and *clr1C* growth in YFP control media an in various stress media: Minimal Media (nutrient limitation), YDP with 1 M NaCl (salt stress), YPD with 50 μg/mL Congo Red or YPD with 40 μg/mL Calcofluor White (cell wall perturbations) or 1 M of Sorbitol (osmotic stress). The experiment was repeated 2 times with similar results. Plates were spot-inoculated with 20 μl of decimated dilution from 10^7^ microconidia/ml.

**Fig. S3.**
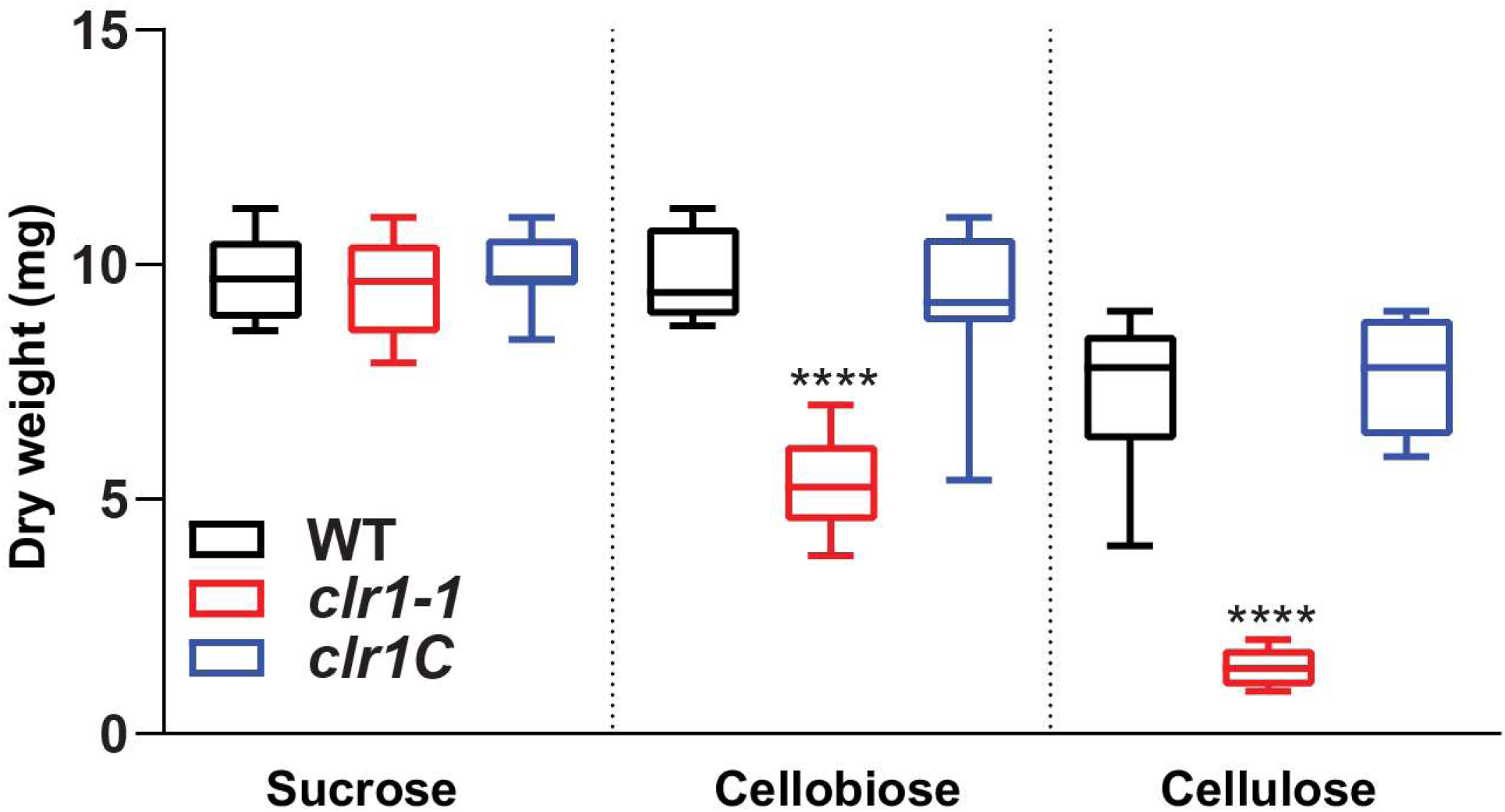
The lack of CLR1 impairs Fo growth in different carbon sources. Growth WT, *clr1-1* and *clr1C* on sucrose 0.5% or cellobiose 0.5% for 3three days, or on cellulose 0.5% for 7 days measured as dry weight (mg). Shown are the box plots: centerlines show the medians; box limits indicate the 25^th^ and 75^th^ percentiles; whiskers extend to the minimum and maximum, N=9. Asterisk indicated differences relative to WT. Welch’s unpaired t-test; **** p-value ≤ 0.0001.

**Fig S4.**
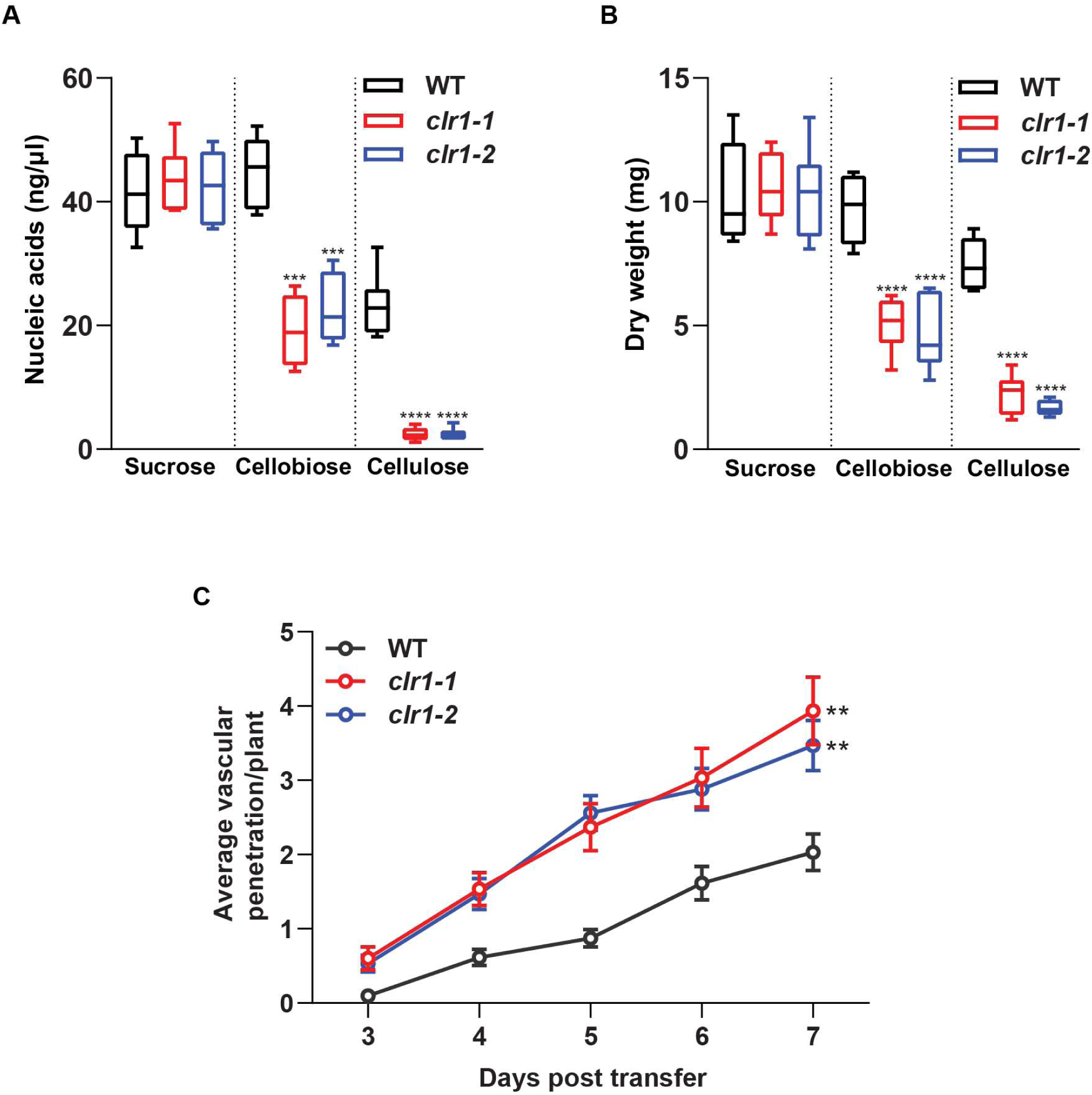
A second independent CLR1 deletion mutant, *clr1-2*, displayed the same phenotype than *clr1-1*. **(A)** and **(B)** Growth of WT, *clr1-1* and *clr1-2* on sucrose 0.5% or cellobiose 0.5% for 3 days, or on cellulose 0.5% for 7 days measured as dry weight (mg) (A) and nucleic acid concentration (ng/ml) (B). Shown are the box plots: centerlines show the medians; box limits indicate the 25^th^ and 75^th^ percentiles; whiskers extend to the minimum and maximum, N≥5. Welch’s unpaired t-test; asterisks show differences with respect to WT.***p-value≤0.001, ****p-value ≤0.0001. **(C)** Cumulative Arabidopsis root vascular penetration by Fo at different days post-transfer (dpt) to WT, *clr1-1* or *clr1-2* microconidia plates. Values are mean +/- SEM, N≥30 plants from one representative experiment. The experiment was performed 3 times with similar results. RM two-way ANOVA on vascular penetration rate: p<0.0001 (fungal genotype), p<0.0001 (time), p ≤ 0.01 (fungal genotype x time). Asterisk indicated a statistical difference with respect to WT at 7 dpt, Tukey’s multiple comparison test, ** p<0.01.

**Fig. S5.**
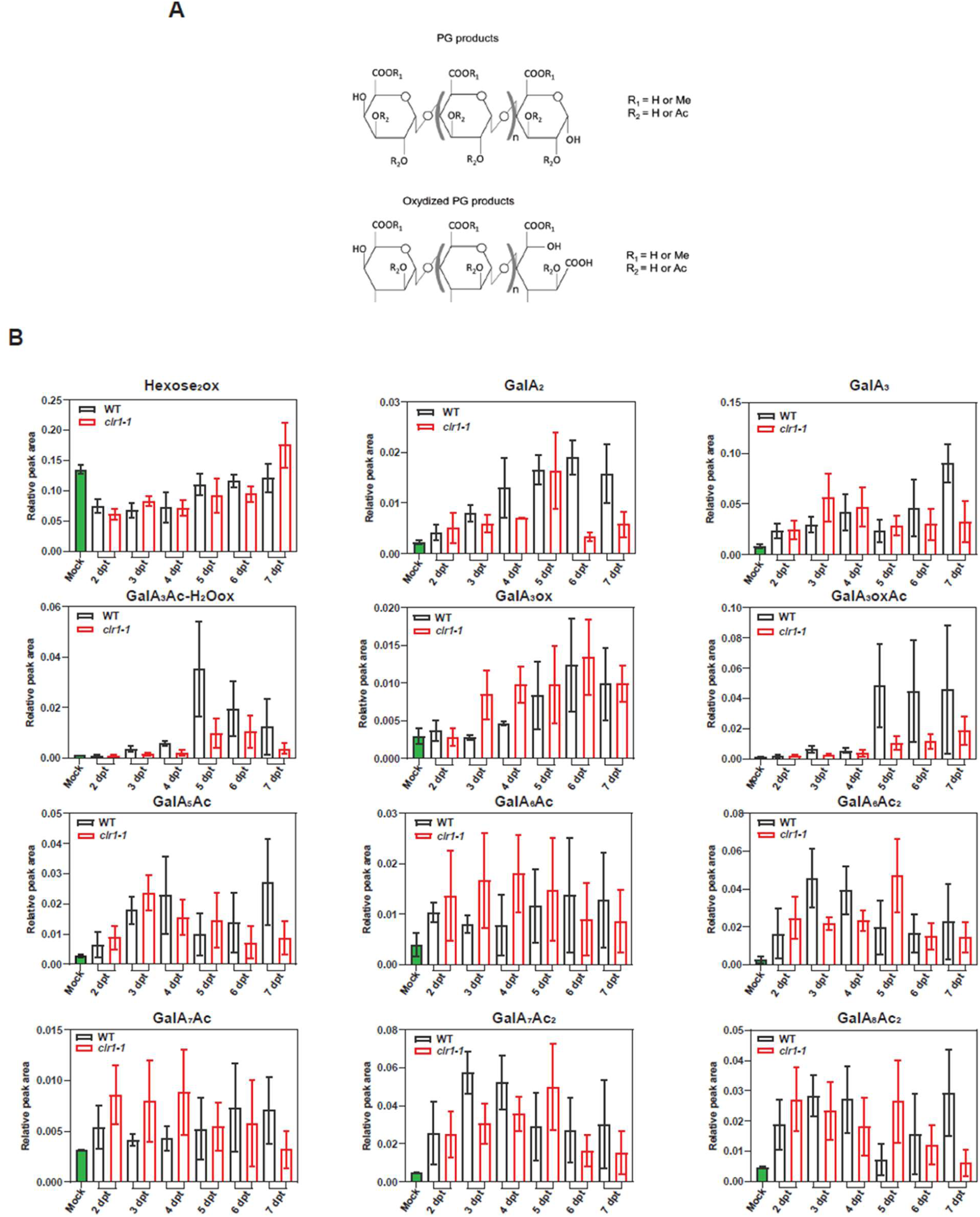
Kinetics of Hexoses and OGs identified in Arabidopsis roots infected with WT or *clr1-1*. **(A)** Examples of the modifications observed in the detected OGs: acetylated or methylated residues and oxidized PG products. **(B)** Kinetics of Hexoses_2_ and OGs identified by HP-SEC-MS in infected roots at different days post transfer (dpt) to WT or *clr1-1* microconidia. Bars represent means +/- SEM from 3 biological replicates (2 for mock samples). No significant differences were observed in the *clr1-1* samples compared to WT ones using two-way ANOVA with LSD Fisher test post hoc comparison at any dpt. OGs are named GalA_x_Ac_y_, indicating the subscript numbers indicate the degree of polymerization (x) and the number of acetylated groups (y). “ox” indicate the presence of oxidized groups, respectively.

**Fig. S6.**
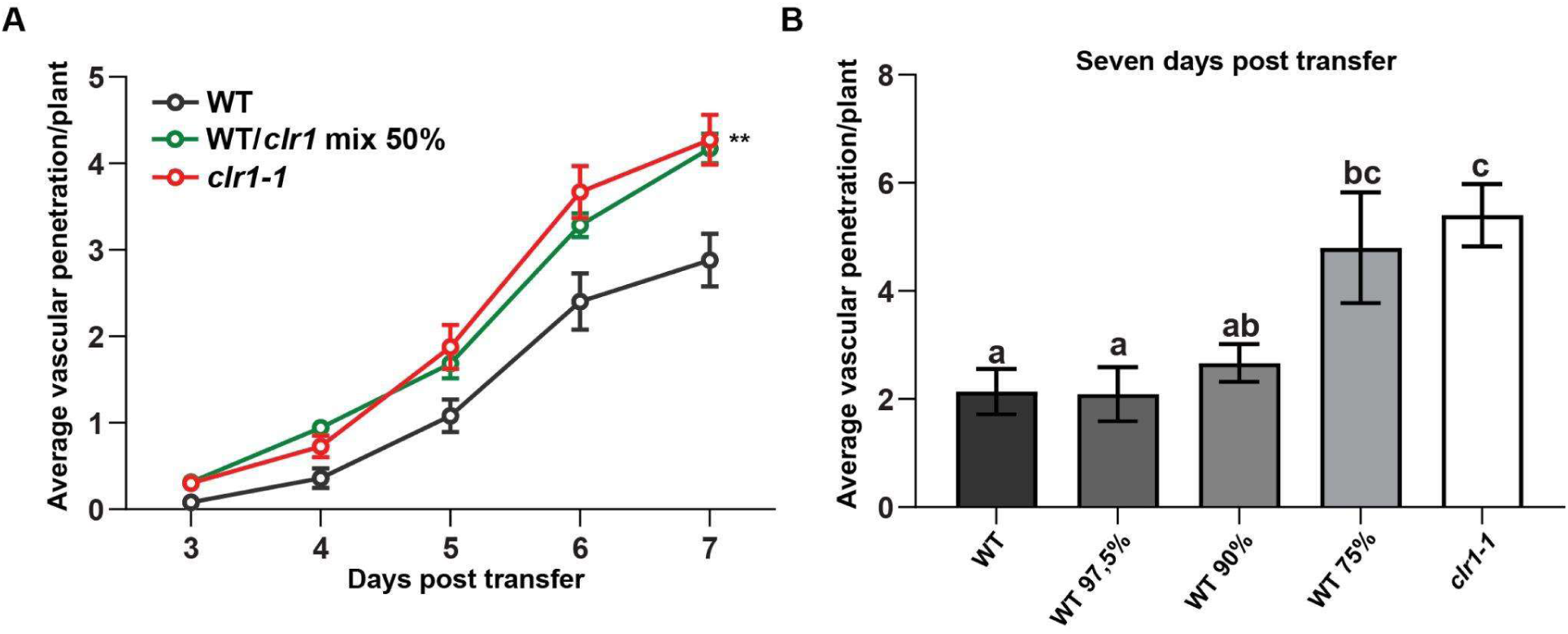
*clr1-1* virulence is not compensated by WT coinfection. **(A)** Cumulative Arabidopsis root vascular penetration by Fo at different days post-transfer (dpt) to plates with WT, *clr1-1* or a 50/50 WT/*clr1-1* microconidia mix. Values are mean +/- SEM, N≥25 plants from one representative experiment. The experiment was performed 3 times with similar results. RM two-way ANOVA on vascular penetration rate: p<0.001 (fungal genotype), p<0.0001 (time), p≤0.01 (fungal genotype x time). Asterisk indicated a statistical difference with respect to WT at 7 dpt. Tukey’s multiple comparisons test, ** p<0.01. **(B)** Root vascular penetration at 7 dpt to plates with WT, *clr1-1* or a mix with 97,5%, 90%, or 75% WT microconidia complemented to 100% with *clr1-1* microconidia. Bars represent means +/- SEM, N≥22, from one independent experiment. The experiment was performed 2 times with similar results. One-way Anova was performed and Tukey’s multiple comparisons test; letters indicate statistical difference between samples; p<0.05.

**Fig. S7.**
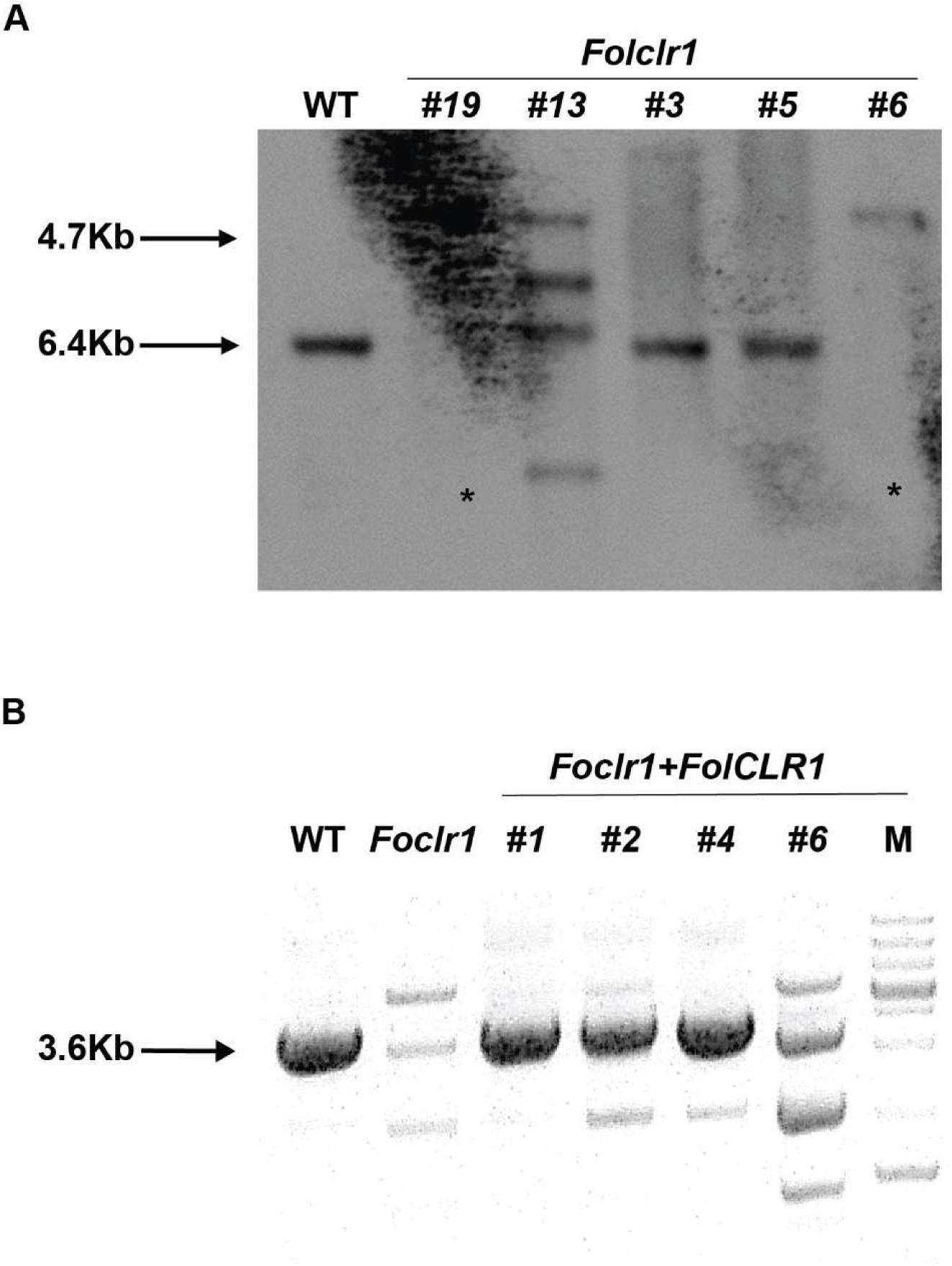
Identification of Fol4287 *clr1* and *clr1C* mutants. **(A)**. Southern blot hybridization analysis of the WT and of five indicated *clr1* transformants. Genomic DNA was digested with the *HindIII* restriction enzyme and hybridized with a probe obtained by using the primer pair CLR1TERFOR and CLR1TERREV. 6.4 Kb indicates WT fragment whereas 4.7 the cassette insertion. Transformants marked with an asterisk indicate strains where targeted replacement of the indicated gene by homologous integration of a single copy of the Neo^r^ resistance cassette occurred. **(B)** PCR screening to identify *clr1C* mutant strains. Genomic DNA of the indicated fungal strains was amplified by using the primer pair CLR1PROMFORNEST and CLR1revgene. The expected PCR fragment for *clr1C* mutants is of 3600 bp.

**Table S1.**
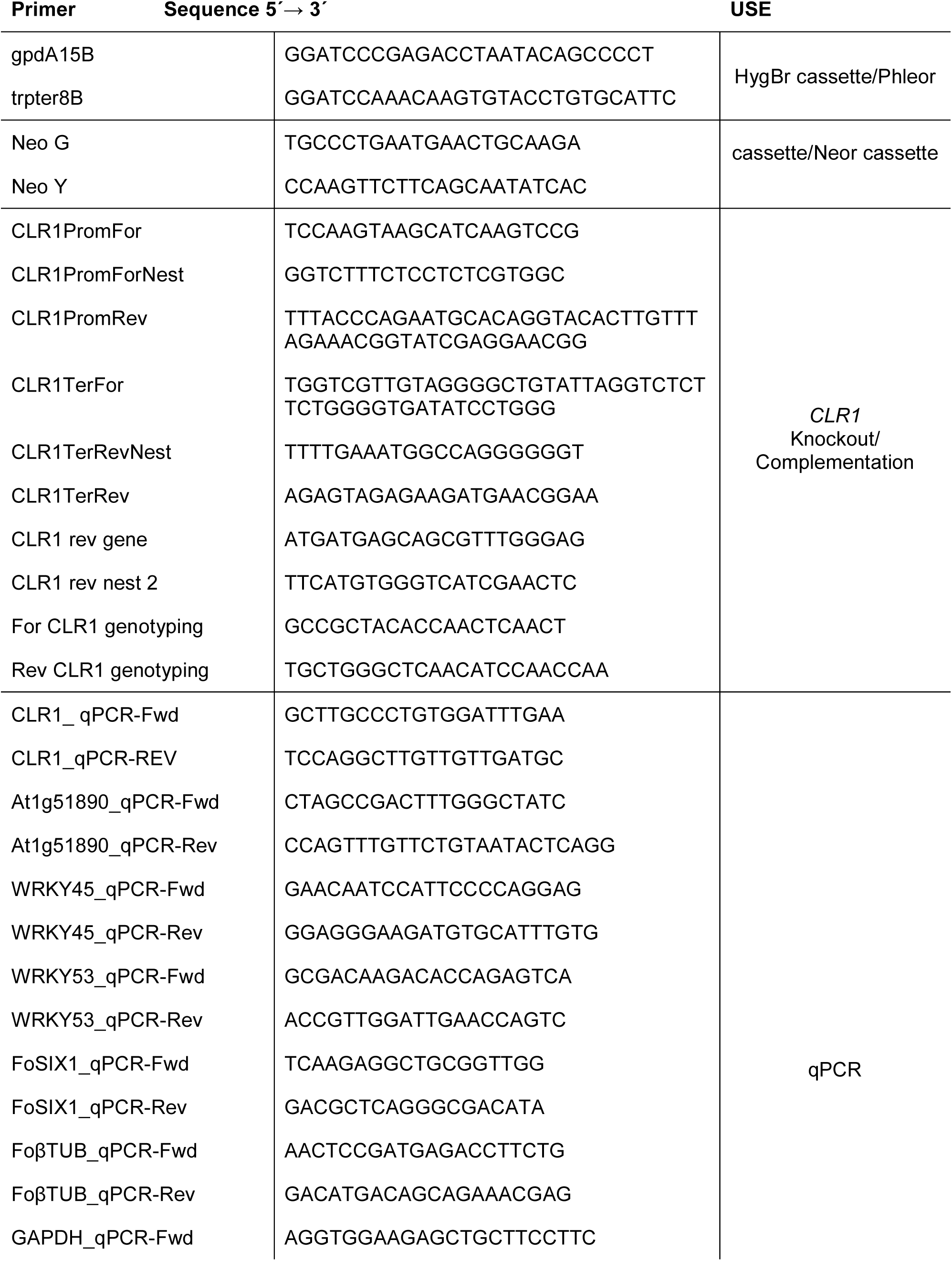
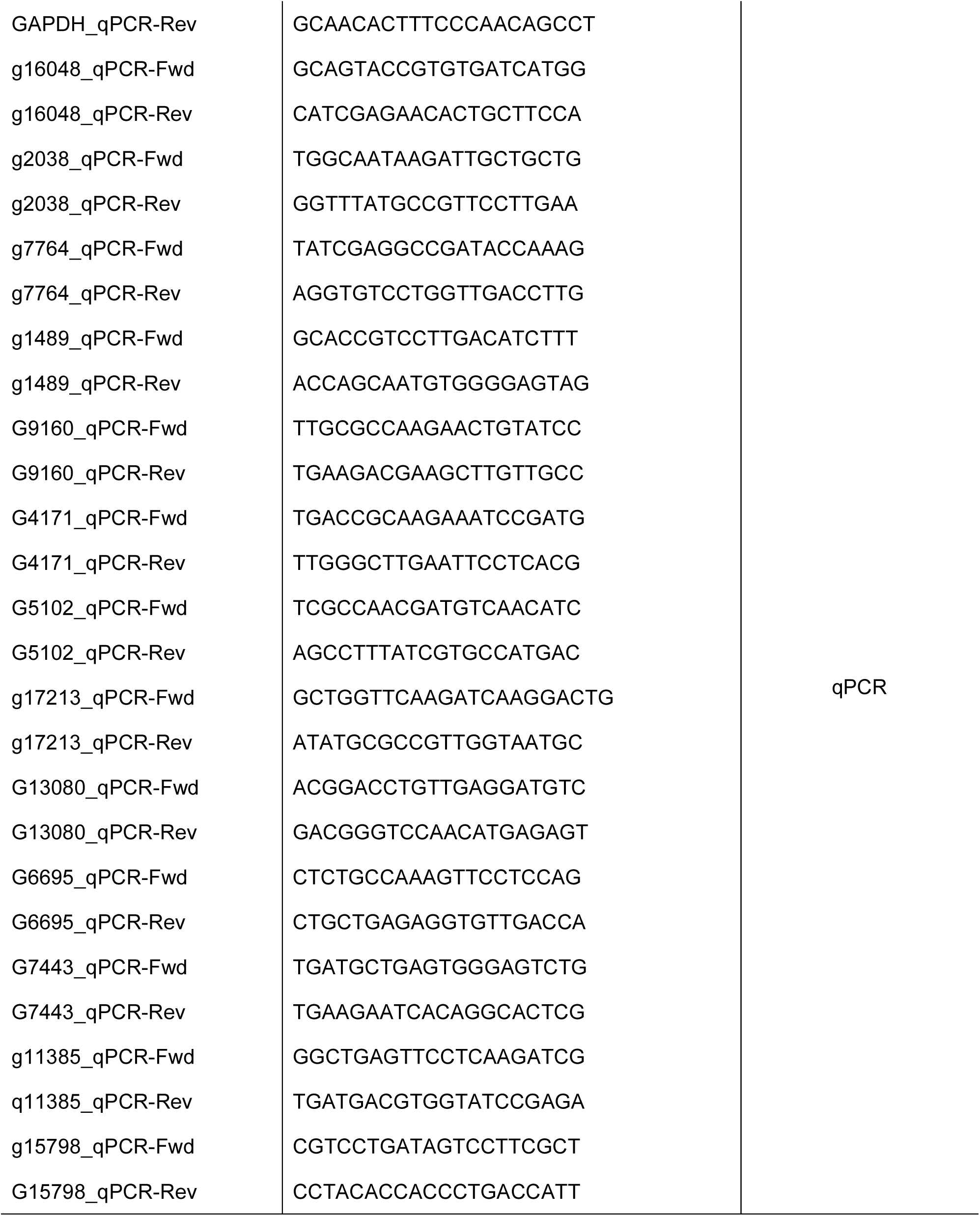
Primers used in this study.

